# Spiperone targets HBV cccDNA via ER stress–induced innate immune activation and epigenetic silencing

**DOI:** 10.64898/2026.03.31.715751

**Authors:** Junghwa Jang, Ziyun Kim, Eunseo Kim, Jisung Park, Yu-Min Choi, Bum-Joon Kim

## Abstract

Chronic hepatitis B persists due to the stability of nuclear covalently closed circular DNA (cccDNA), which maintains viral transcription despite prolonged antiviral therapy, highlighting the need for strategies that suppress cccDNA via host-targeted mechanisms. Here, we identify Spiperone, a clinically approved compound, as a repurposed anti-HBV candidate with strong translational potential. Spiperone robustly reduced HBsAg, HBeAg, viral DNA, and pgRNA across HepG2.2.15, HBV-infected HepG2-NTCP-C4 and HepaRG cells, and multiple in vivo models, including HBV transgenic, hydrodynamic injection, and AAV- HBV1.04× models. Notably, intrahepatic cccDNA was significantly diminished. In combination, Spiperone potentiated tenofovir activity, exhibiting synergistic effects, while both intraperitoneal and oral administration reduced antigenemia and viremia.

Mechanistically, Spiperone activated the PERK–eIF2α–ATF4 arm of the ER stress response, coupled with mitochondrial perturbation and cytosolic release of oxidized mitochondrial DNA, leading to activation of IFI16–STING–IRF3 signaling. This cascade induced type I interferon (IFN-I) and interferon-stimulated genes. ChIP–qPCR further demonstrated reduced enrichment of activating histone marks on cccDNA, consistent with transcriptional repression.

Collectively, these findings position Spiperone as a host-directed antiviral that converges ER stress–linked innate immunity and epigenetic repression to suppress cccDNA, supporting its advancement in combination strategies toward a functional cure for chronic HBV infection.

## 1. Introduction

Chronic hepatitis B virus (HBV) infection is a major causative factor in the pathogenesis of severe hepatic disorders, including cirrhosis and hepatocellular carcinoma (HCC)^1–3^. HBV-related complications are still estimated to account for approximately 1.1 million deaths annually, and about 15–25% of individuals with chronic HBV infection ultimately succumb to cirrhosis or HCC ^4, 5^. Despite the availability of preventive vaccines and effective therapies ^6, 7^, no treatment has yet achieved a functional cure for HBV. The major reason for this limitation is the persistence of covalently closed circular DNA (cccDNA), which remains a key obstacle to cure.

In infected hepatocytes, HBV establishes a cccDNA minichromosome within the nucleus, which serves as the transcriptional template for all viral pregenomic RNAs (pgRNAs) and messenger RNAs (mRNAs)^8, 9^. Despite the potent suppression of viremia achieved through the administration of nucleos(t)ide analogs, cccDNA remains a persistent reservoir even after extended treatment periods ^10^. Interferon-α (IFN-α)-based therapies also show limited efficacy in achieving sustained cccDNA clearance^11–15^. In addition, cccDNA has a long half-life, and the cccDNA pool in infected hepatocytes can be replenished through continuous cycles of infection and intracellular amplification^16, 17^. Consistent with these characteristics, HBV reactivation has been detected in individuals with sustained loss of HBsAg following treatments such as chemotherapy, immunosuppressive therapy, or hematopoietic stem cell transplantation. Thus, effective strategies for HBV functional cure require not only suppression of viral replication but also elimination or durable silencing of cccDNA^18, 19^.

To overcome this obstacle, diverse antiviral strategies targeting cccDNA are currently under investigation. In particular, approaches that block viral entry or nucleocapsid recycling can suppress *de novo* cccDNA formation but have limited effects on the established cccDNA pool^20, 21^. In contrast, genome-editing strategies can directly target pre-existing cccDNA^22–24^. However, their clinical application is limited by inefficient delivery and safety concerns, including imperfect repair, chromosomal rearrangements, and off-target effects^25, 26^. Therefore, alternative therapeutic strategies that can effectively eliminate or functionally silence persistent cccDNA are urgently needed.

Given these limitations, host-directed antiviral strategies have attracted increasing attention as a promising alternative for achieving HBV functional cure. In particular, hepatocyte-intrinsic stress responses, innate immune signaling pathways, and epigenetic regulation of the cccDNA minichromosome are recognized as key determinants of HBV persistence and transcriptional activity. Among these, IFN-I signaling plays a central role in antiviral defense by suppressing HBV replication and restricting the transcriptional activity of cccDNA^27^. Moreover, previous studies have shown that IFN-I-associated pathways also contribute to the reduction or degradation of cccDNA^28, 29^. Therefore, pharmacologic agents capable of activating host antiviral defense programs, particularly IFN-I-associated signaling pathways, may represent a clinically feasible strategy to suppress HBV replication while promoting the reduction or functional silencing of persistent cccDNA.

Drug repurposing involves exploring new therapeutic uses for drugs that have already been approved for other indications^30^. This approach enables the identification of potential treatments for different diseases, offering considerable advantages over traditional drug discovery in terms of cost-efficiency and accelerated development timelines ^31, 32^. Spiperone, a selective antagonist targeting the dopamine D2 receptor and the 5-HT1A/2A receptors, which are G-protein coupled receptors (GPCRs), was approved for the treatment of schizophrenia in Japan in 1969 ^33–36^. Beyond its primary indication, Spiperone has been investigated for potential therapeutic applications in a variety of diseases, including neurodegenerative diseases and cancers^37–39^. Given its well-established safety profile, Spiperone is a promising candidate for drug repurposing. In this study, we aimed to repurpose Spiperone as a host-directed anti-HBV agent capable of reducing and functionally silencing cccDNA and to elucidate the underlying mechanism.

## 2. Results

### 2.1 Spiperone effectively suppresses HBV antigen, DNA, and cccDNA levels in HepG2.2.15, HBV-infected HepG2-NTCP-C4 and HepaRG cells

To identify potential anti-HBV candidates, we screened 500 compounds from a Korea Chemical Bank library consisting of phase I–III clinical drugs and approved compounds. The stable HBV-producing HepG2.2.15 cell system was used to assess both HBsAg secretion and cytotoxicity (Supplementary Fig. S1A and B). Ten compounds were selected based on their initial suppression of HBsAg and further evaluated at increasing concentrations (5, 10, and 20 μM) to assess dose-dependent effects on HBsAg reduction and cytotoxicity (Supplementary Fig. S2A and B). Among these compounds, Spiperone was identified as the most effective in reducing HBsAg levels while maintaining acceptable cytotoxicity.

To evaluate its effect on HBV replication, HepG2.2.15 cells were treated with Spiperone for 48 h. Compared with PBS or tenofovir disoproxil fumarate (TDF), Spiperone significantly reduced HBsAg, HBeAg and markedly decreased HBV DNA levels (Fig. 1A). The dose-response curve shows that Spiperone decreased the HBsAg level, with an /C_SO_ of approximately 1.3 μM (Supplementary Fig. S3). Analysis of pgRNA levels revealed that Spiperone treatment led to a marked decrease in the level of pgRNA, which is regarded as a surrogate clinical biomarker of intrahepatic cccDNA activity^40, 41^. These findings indicate that Spiperone suppresses cccDNA transcriptional activity (Fig. 1B). Consistently, Spiperone treatment reduced HBV capsid and hepatitis B core antigen (HBcAg) level in a dose-dependent manner, as confirmed by western blot (Fig. 1C and Supplementary Fig. S4) and confocal microscopy (Fig. 1D). Following HBV infection of HepG2-NTCP-C4 cells, treatment for 48 h with Spiperone reduced HBsAg, extracellular HBV DNA, and intracellular pgRNA levels more effectively than PBS or TDF consistent with the results observed in HepG2.2.15 cells (Fig. 1E). We next assessed the antiviral activity of Spiperone in differentiated HepaRG cells, which permit HBV entry and cccDNA formation. Spiperone similarly reduced HBsAg secretion and lowered both viral DNA and pgRNA levels in this system (Fig. 1F). Based on the effects of Spiperone on viral replication and its impact on cccDNA transcriptional activity, we next examined whether Spiperone directly affects cccDNA levels. The level of cccDNA was quantified using a cccDNA-selective primer set. Spiperone decreased the cccDNA level in HepG2.2.15 (*p* < 0.01 vs PBS; *p* < 0.001 vs TDF), HepG2-NTCP-C4 (*p* < 0.01 vs PBS; *p* < 0.01 vs TDF), and HepaRG cells (*p* < 0.01 vs PBS; *p* < 0.05 vs TDF) (Fig. 1G) Furthermore, PCR analysis of extracted DNA demonstrated that Spiperone markedly reduced cccDNA levels, as evidenced by the diminished cccDNA-specific band, while non-selective primers failed to produce detectable bands in all groups (Fig. 1H). Southern blot analysis further confirmed the reduced cccDNA band in Spiperone-treated cells (Fig. 1I). To validate this effect, IFN-α, a known inhibitor of HBV replication and cccDNA[², was used as a positive control. Spiperone suppressed HBsAg secretion and reduced cccDNA levels (*p* < 0.001 vs PBS; *p* < 0.05 vs TDF) more efficiently than IFN-α (Fig. 1J, K). Consistently, PCR analysis confirmed a marked reduction in cccDNA following Spiperone treatment (Fig. 1L). Given that combination antiviral therapy is a widely adopted strategy for enhancing antiviral efficacy and suppressing the emergence of resistance, we further investigated the combined effects of Spiperone and TDF, a first-line therapy for chronic HBV infection with a well-established safety profile.^42^ To evaluate the potential synergy between Spiperone and TDF, drug combination responses were assessed using the SynergyFinder version 3.0 with the ZIP reference model. The analysis yielded a synergy score greater than 5, and the results were visualized with two- and three-dimensional synergy maps (Fig. 1M). Consistent with these findings, combination treatment suppressed HBsAg more effectively than either monotherapy, as shown in Supplementary Fig. S5. Taken together, these results demonstrate that Spiperone reduces HBV antigens, viral DNA, and pgRNA levels and decreases cccDNA in both stable replication systems and HBV infection models.

**Fig. 1.**
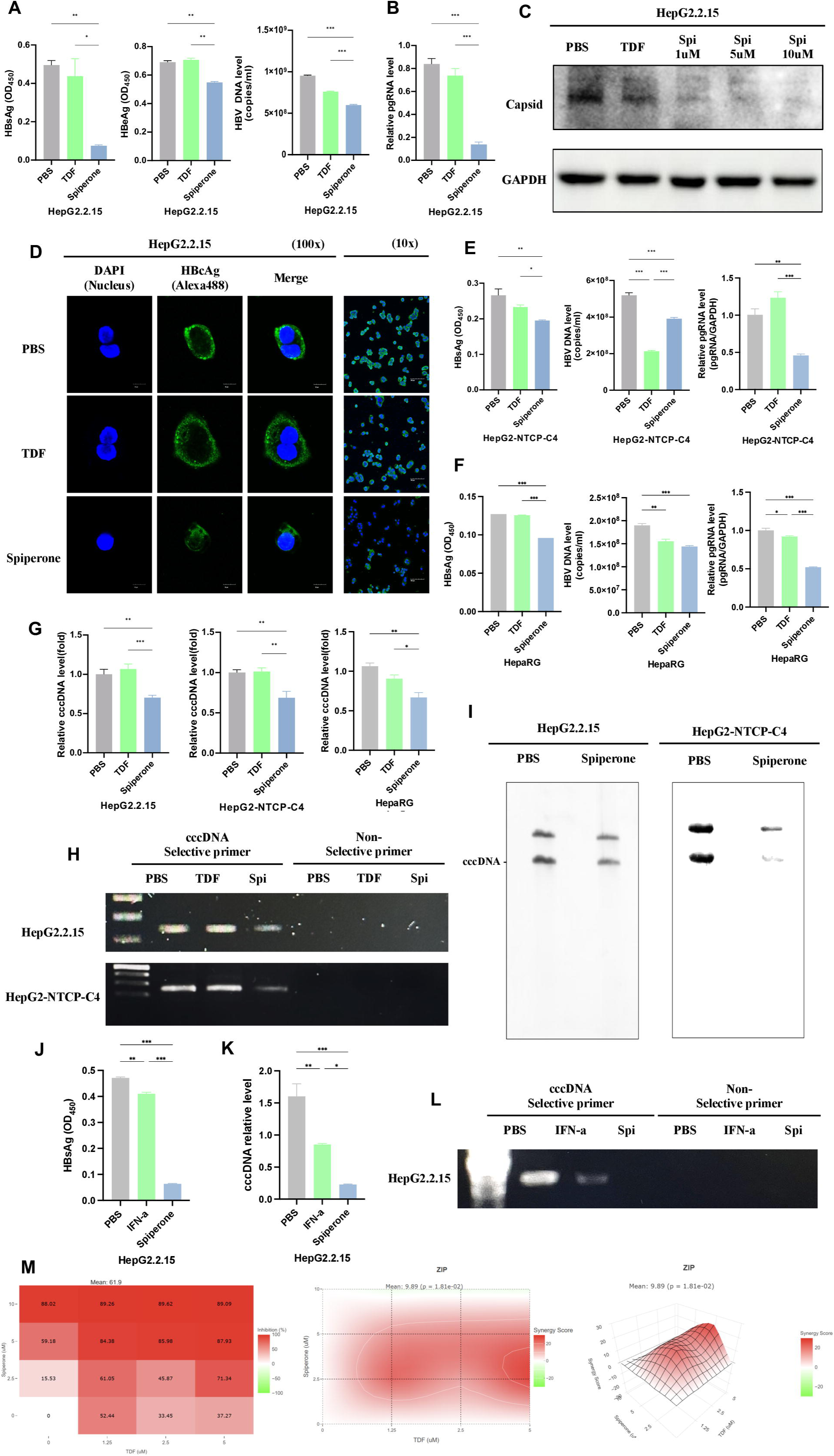
Spiperone reduces HBV antigen, DNA, and cccDNA in HepG2.2.15, HepG2-NTCP-C4, and HepaRG cells. (A) HBsAg, HBeAg and HBV viral DNA levels were quantified by ELISA and qPCR. (B) pgRNA level was measured by qPCR. (C and D) Capsid protein level was observed by western blotting (C) and HBcAg expression (Alexa488) was observed by confocal microscopy (D) in HepG2.2.15. The representative images are shown. (E and F) The level of HBsAg, HBV DNA and pgRNA in HBV-infected HepG2-NTCP-C4 cell (E) and HepaRG cell (F) were measured. (G) The cccDNA levels were measured by qPCR in HepG2.2.15, HBV-infected HepG2-NTCP-C4 and HepaRG cells. (H) Gel electrophoresis images showing cccDNA using cccDNA selective primer or non-selective primer in PBS-, TDF-, or Spiperone-treated cells. (I) Southern blot images showing cccDNA band in HepG2.2.15 and HepG2-NTCP-C4 cells treated with PBS or Spiperone. (J-L) HepG2.2.15 cells were treated with PBS, IFN-α, or Spiperone. The level of HBsAg was measured by ELISA (J). cccDNA were evaluated by qPCR (K) and gel electrophoresis using cccDNA selective primer or non-selective primer (L). (M) The synergistic effect of TDF and Spiperone was shown in two- and three-dimensional synergy maps. The significance of differences is indicated as *p < 0.05, **p < 0.01, and ***p < 0.001. HBV, Hepatitis B virus; HBsAg, HBV surface antigen; HBeAg, HBV e antigen; HBcAg, HBV core antigen; qPCR, quantitative polymerase chain reaction; pgRNA, pregenomic RNA; cccDNA, covalently closed circular DNA; TDF, tenofovir disoproxil fumarate.

### 2.2 Antiviral effects of Spiperone in HBV transgenic mice following intraperitoneal and oral administration

To evaluate the *in vivo* antiviral efficacy of Spiperone, HBV transgenic (TG) mice were administered daily intraperitoneal injections of PBS (*n* = 5 mice), TDF (500 μg per mouse; *n* = 5 mice), or Spiperone (1.5 mg per kg; *n* = 5 mice) for eight weeks (Fig. 2A). Serum samples were collected, and HBV antigen concentrations were measured by ELISA. Spiperone treatment resulted in a significant decrease in the concentrations of both HBsAg and HBeAg (Fig. 2B, C), with a consistent reduction in the HBsAg concentration observed at weeks 2, 3 and 4 (Supplementary Fig. S6). Furthermore, serum HBV DNA concentrations were reduced in both the TDF- and Spiperone-treated groups, although the decrease was more substantial with Spiperone treatment (Fig. 2D). To further validate the effect of Spiperone at the tissue level, HBcAg expression was evaluated by immunohistochemistry (IHC), and HBsAg expression was evaluated by confocal microscopy. Compared with the PBS or TDF groups, Spiperone treatment resulted in a clear reduction in hepatic HBcAg staining (Fig. 2E). Consistently, HBsAg expression was markedly lower in the group treated with Spiperone, supporting its inhibitory effect on antigen expression (Fig. 2F). Chronic HBV infection typically requires prolonged antiviral therapy. Given that Spiperone has been used as an oral antipsychotic agent (*Spiropitan*) in Japan, evaluating whether it retains antiviral activity following oral administration is critical for its therapeutic repurposing. To evaluate the effects of oral administration, HBV TG mice were treated by daily gavage with PBS (n = 5), Spiperone (n = 5), or TDF (n = 5) at the same doses used for intraperitoneal injection for eight weeks (Fig. 2G). Consistent with the effects observed following intraperitoneal administration, oral Spiperone treatment significantly reduced serum HBsAg, HBeAg, and HBV DNA levels (Fig. 2H-J). Liver tissues from orally treated mice showed reduced HBcAg and HBsAg staining similar to that observed following intraperitoneal administration (Fig. 2K, L), confirming that Spiperone retains antiviral activity across different routes of administration.

**Fig. 2.**
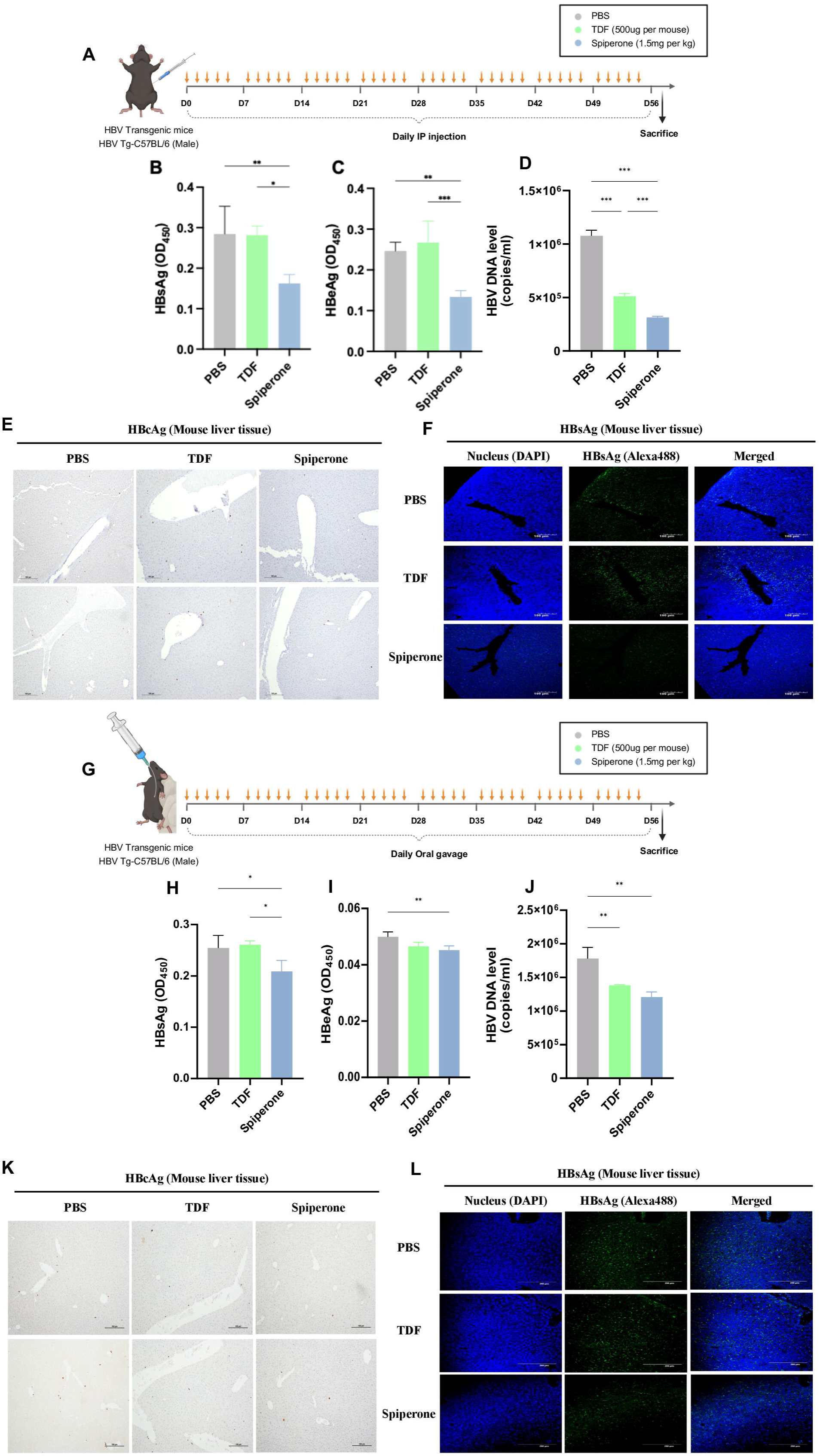
Antiviral effects of Spiperone in HBV transgenic mice following intraperitoneal and oral administration. (A) Schematic illustration of the process for Spiperone administration (IP) in HBV transgenic mice. (male; *n* = 5 per each group) (B-D) The levels of HBsAg (B), HBeAg (C) and HBV DNA (D) in mouse serum samples were assessed by ELISA or qPCR. (E and F) IHC images showing HBcAg expression (E) and confocal images showing HBsAg expression (Alexa Fluor 488) (F) in mouse liver tissue. The representative images are shown. (G) Schematic illustration of the process for Spiperone administration (oral) in HBV TG mice (Male; *n* = 5 per each group) (H-J) The levels of HBsAg (H), HBeAg (I) and HBV DNA (J) in mouse serum samples were assessed by ELISA or qPCR. (K and L) IHC images showing HBcAg expression (K) and confocal images showing HBsAg expression (Alexa Fluor 488) (L) in mouse liver tissue. The representative images are shown. The significance of differences is indicated as *p < 0.05, **p < 0.01, and ***p < 0.001. HBV, Hepatitis B virus; TG mouse, Transgenic mouse; IP, Intraperitoneal injection; ELISA, Enzyme-Linked ImmunoSorbent Assay; IHC, Immunohistochemical.

### 2.3 Spiperone activates the PERK-eIF2**α**-ATF4 axis of the ER stress response in HBV-infected hepatocytes

To elucidate the molecular mechanisms underlying the observed antiviral activity of Spiperone, we performed RNA-sequencing analysis of Spiperone-treated hepatocytes. Pathway enrichment and volcano plot analyses revealed prominent upregulation of ER stress–related responses, suggesting a potential role for ER stress signaling in mediating the antiviral effects of Spiperone (Fig. 3A, B and Supplementary Fig S7). RNA-sequencing revealed transcriptional signatures characteristic of ER stress following Spiperone exposure (Fig. 3C). Consistent with these findings, qPCR analysis in HBV-infected HepG2-NTCP-C4 and HepG2.2.15 cells showed increased expression of PKR-like ER kinase (PERK), activating transcription factor 4 (ATF4), activating transcription factor 6 (ATF6), and X-box binding protein 1 (XBP1), with the most pronounced induction observed in the PERK–eIF2α–ATF4 branch (Fig. 3D, E). This pattern was recapitulated in differentiated HepaRG cells, where Spiperone increased PERK and ATF4 mRNA levels (Fig. 3F), further supporting activation of the PERK-dependent branch of the ER stress response in a physiologically relevant hepatocyte model. At the protein level, western blot analysis showed increased levels of phosphorylated PERK, phosphorylated eIF2α, and ATF4 in HepG2.2.15 cells (Fig. 3G). Immunofluorescence staining further confirmed enhanced nuclear accumulation of ATF4, supporting activation of this pathway (Fig. 3H). This effect was observed in a concentration-dependent manner (Supplementary Fig. S8). IHC analysis of liver tissue from HBV TG mice revealed that compared with PBS or TDF, Spiperone treatment increased hepatic p-PERK and ATF4 levels (Fig. 3I, J). Taken together, these data indicate that Spiperone selectively activates the PERK–eIF2α–ATF4 axis of the ER stress response in HBV-infected hepatocytes.

**Fig. 3.**
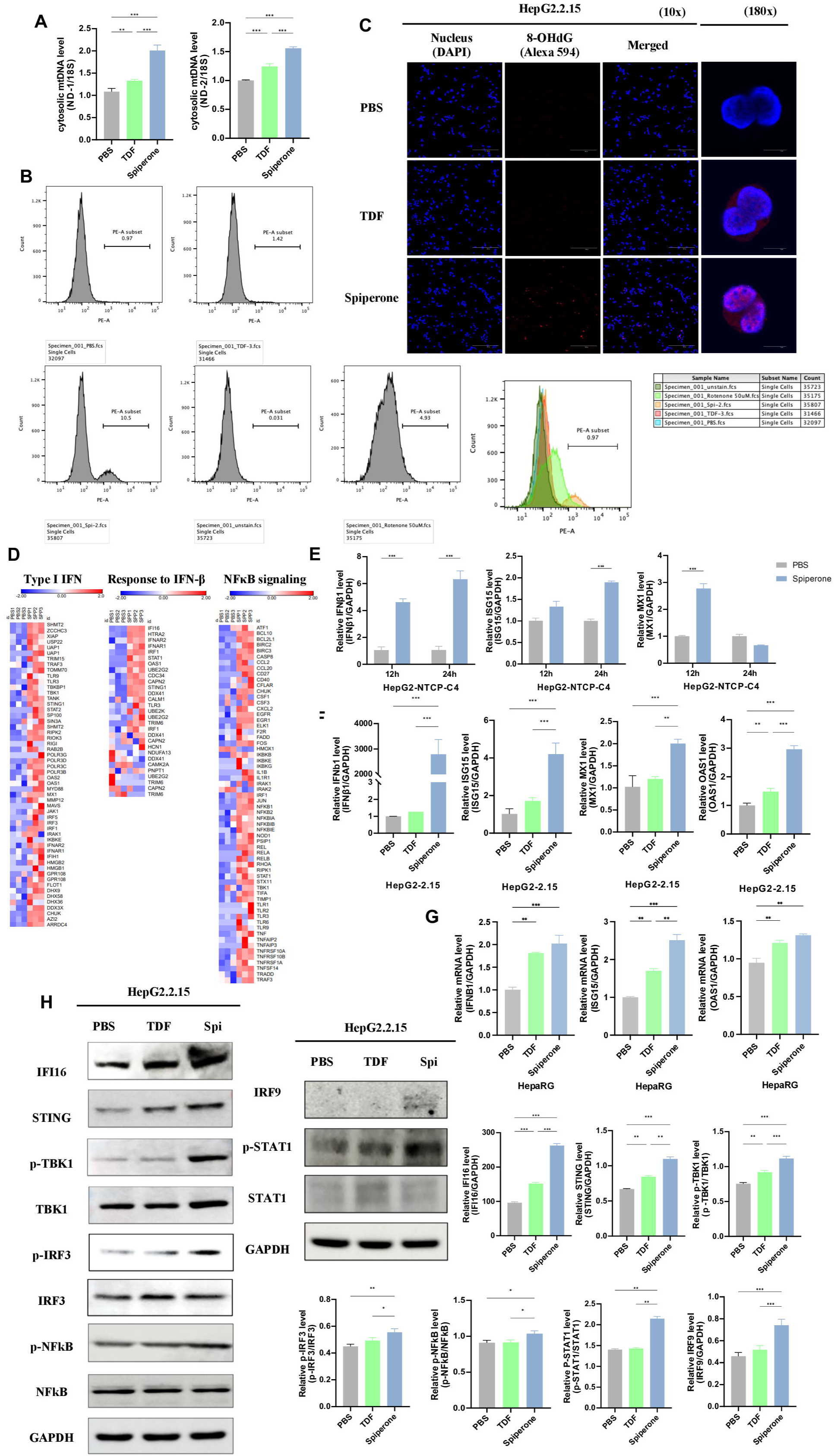
Spiperone activates the PERK-eIF2α-ATF4 axis of the ER Stress response in HBV-infected hepatocytes. (A-C) RNA-seq analysis was performed in HBV-infected HepG2-NTCP-C4 cell. The bar plot (A) shows diverse upregulated pathways. Volcano plot (B) and heatmap (C) show upregulated responses to ER stress. The heatmap visualizes relative expression patterns using gene-wise min–max–normalized expression values. (D-F) The relative mRNA level of ER stress-related genes in HepG2-NTCP-C4 (D), HepG2.2.15 (E) and HepaRG cells (F). (G) Western blot showing ER stress-related proteins. The intensities were quantified using ImageJ and normalized to GAPDH (or non-phosphorylated form). (H) Confocal images showing ATF4 nuclear accumulation (Alexa Fluor 488) in HepG2.2.15 cell (20X and 100X magnification). (I and J) Representative IHC images of p-PERK (I) and ATF4 (J) in liver tissue from TG mice administered with PBS, TDF or Spiperone. The significance of differences is indicated as *p < 0.05, **p < 0.01, and ***p < 0.001. HBV, Hepatitis B virus; ER, Endoplasmic Reticulum; IHC, Immunohistochemical.

### 2.4 Spiperone promotes mitochondrial stress and activates DNA-sensing pathways to induce IFN-I responses

Given that Spiperone increases ER stress in HBV-infected cells and that ER stress is known to elicit mitochondrial dysfunction and stress responses ^43, 44^, we next examined whether Spiperone triggers mitochondrial stress. RT–qPCR analysis of the cytosolic fraction revealed significant increases in the levels of multiple mitochondrial DNA markers in Spiperone treated cells, indicating the release of mtDNA into the cytoplasm (Fig. 4A, Supplementary Fig. S9). Flow cytometric analysis with MitoSOX staining further showed a marked increase in the proportion of ROS-accumulating cells following Spiperone treatment, consistent with the elevated mitochondrial superoxide levels (Fig. 4B). In addition, confocal microscopy visualization of 8-hydroxy-2′-deoxyguanosine (8-OHdG), a marker of oxidized DNA, revealed a strong signal in Spiperone-treated cells (Fig. 4C). This finding suggests that the released mtDNA was present in a damaged and oxidized form, and its level increased in a concentration-dependent manner (Supplementary Fig. S10).

**Fig. 4.**
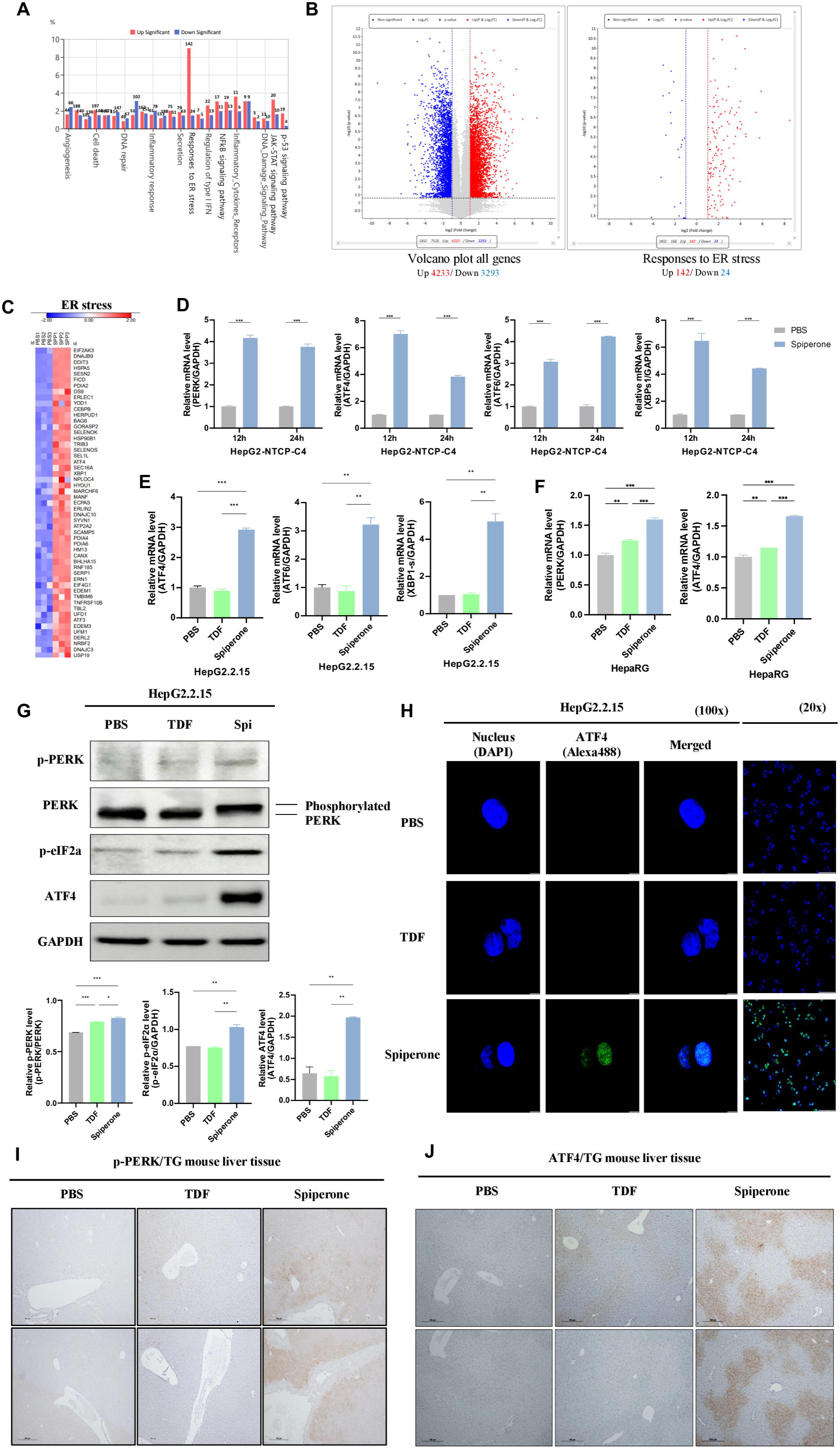
Spiperone promotes mitochondrial stress and activates DNA-sensing pathways to induce IFN-I responses. (A) The level of cytosolic mitochondrial DNA (ND-1 or ND-2) was measured and normalized to 18S rRNA (housekeeping gene). (B) Flow cytometry with MitoSOX staining was performed in response to TDF or Spiperone treatment. (C) Confocal images showing 8-OHdG release (Alexa Fluor 594) in HepG2.2.15 cell (10X and 180X magnification). The representative images are shown. (D) Heatmap of the results of the RNA-seq analysis showing the upregulation of IFN-I signaling, response to IFN-β, and NFkB signaling pathways. The heatmap visualizes relative expression patterns using gene-wise min–max–normalized expression values. (E-G) The relative mRNA level of IFN-I-related genes in HepG2-NTCP-C4 (E), HepG2.2.15 (F) and HepaRG cells (G). (H) Western blot showing IFN-I signaling proteins. The intensities were quantified using ImageJ and normalized to GAPDH (or non-phosphorylated form). The significance of differences is indicated as *p < 0.05, **p < 0.01, and ***p < 0.001. IFN, Interferone; 8-OHdG, 8-Hydroxy-2’-deoxyguanosine.

To investigate the downstream impact of this mitochondrial stress, RNA-sequencing data were analyzed. The clustering heatmap revealed the transcriptional activation of IFN-I signaling, IFN-β responses, and nuclear factor kappa-light-chain-enhancer of activated B cells (NF-kB) pathways following Spiperone exposure (Fig. 4D). Notably, pathway analysis identified IFI16, an innate immune DNA sensor, as a convergent and consistently upregulated gene across multiple enriched categories, including pathways related to double-stranded DNA binding, innate immune activation, and cellular responses to interferon-β (Supplementary Fig. S11).^45^ Consistent with this observation, STING, a key downstream mediator of IFI16 signaling, also exhibited increased mRNA expression in Spiperone-treated cells (p < 0.001 vs PBS), as shown in the supplementary data (Supplementary Fig. S12). These findings suggest that IFI16 may act as a key link connecting DNA sensing, cytosolic mtDNA, and innate immune activation. Consistent with this observation, qPCR validation confirmed significant upregulation of representative ISGs, including MX dynamin-like GTPase 1 (MX1), Interferon-stimulated gene 15 (ISG15), IFNβ1, and 2’-5’-oligoadenylate synthetase 1 (OAS1), in both HepG2-NTCP-C4 and HepG2.2.15 cells (Fig. 4E, F). Comparable increases in IFNβ1, ISG15 and OAS1 mRNA levels were also observed in differentiated HepaRG cells (Fig. 4G), reinforcing the activation of IFN-I signaling by Spiperone in a physiologically relevant hepatocyte model. Consistent with the transcriptomic data, western blot analysis in Spiperone-treated HepG2.2.15 cells further demonstrated increased levels of highlighted DNA sensing and innate immune signaling proteins, including IFI16, STING, phosphorylated TANK-binding kinase 1 (TBK1), phosphorylated interferon regulatory factor 3 (IRF3), phosphorylated signal transducer and activator of transcription 1 (STAT1) (Fig. 4H). Importantly, these protein-level changes were consistent with the RNA-seq data (Fig. 4D), indicating coordinated activation of an IFI16–STING–dependent innate immune response. Collectively, our data support a model in which Spiperone-induced ER stress drives mitochondrial DNA release, leading to IFI16-mediated sensing and activation of downstream IFN-I signaling in HBV-infected hepatocytes.

### 2.5 The antiviral effect of Spiperone depends on ER stress–driven DNA sensing and IFN-I signaling

To determine whether the antiviral activity of Spiperone depends on ER stress and DNA-sensing pathways, we performed functional knockdown experiments. Neither ATF6 inhibition nor inositol-requiring enzyme 1 (IRE1) inhibition altered the ability of Spiperone to suppress HBsAg secretion (Supplementary Fig. S13), whereas silencing of eIF2α markedly impaired its ability to reduce the HBV DNA level, underscoring the central role of the PERK–eIF2α–ATF4 branch. (Fig. 5A) Similarly, silencing of IFI16 attenuated the ability of Spiperone to lower the level of HBV DNA compared with scramble controls (Fig. 5B). Luciferase reporter assays using the IFN-β promoter showed that Spiperone-induced transcriptional activity was significantly attenuated upon knockdown of eIF2α, IFI16, or STING, demonstrating that this effect is dependent on these factors. (Fig. 5C-E).

**Fig. 5.**
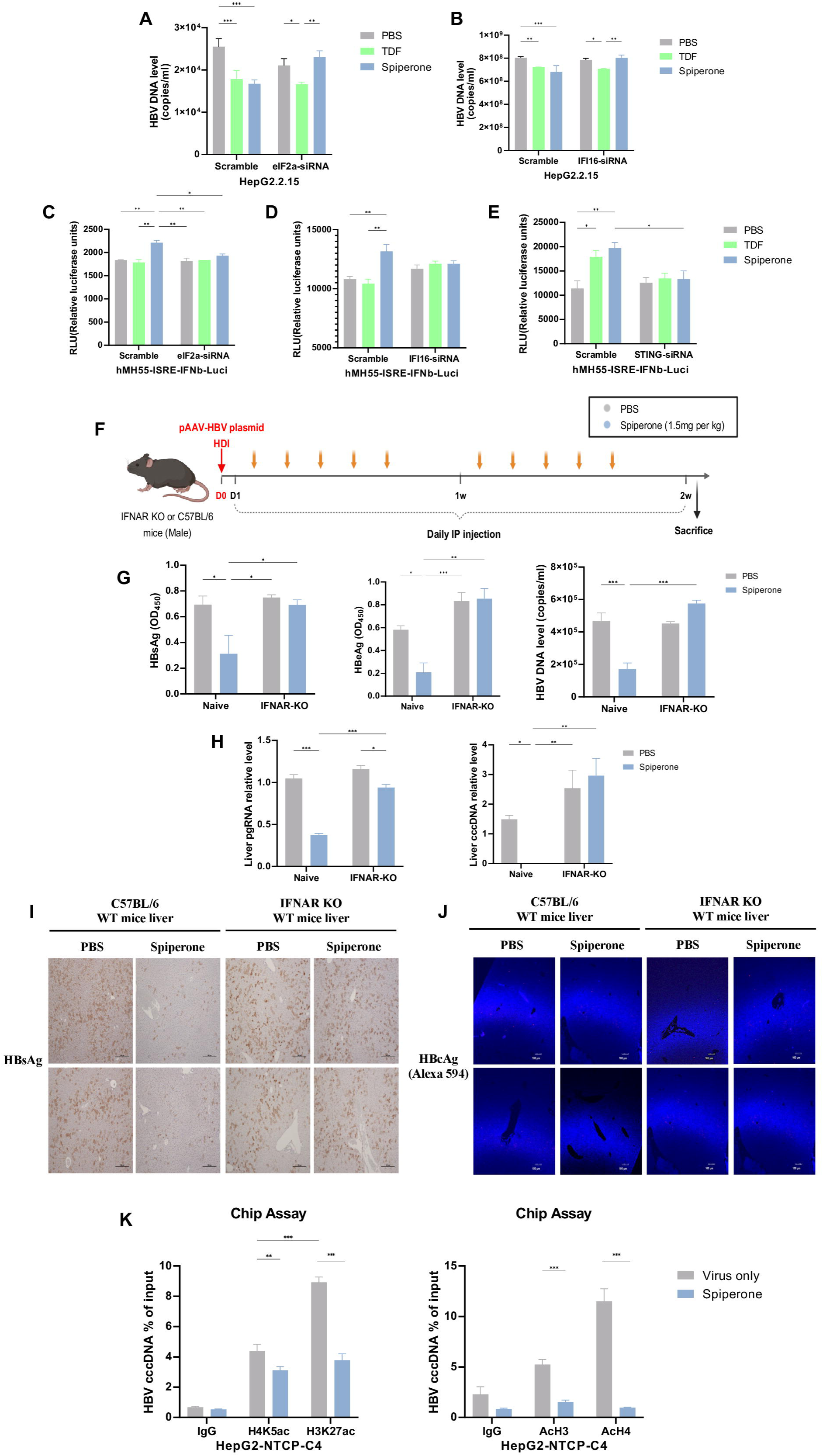
Spiperone’s antiviral effect lies on ER stress-driven DNA sensing, IFN signaling, and cccDNA epigenetic modulation. (A and B) HBV DNA levels after eIF2α (A) or IFI16 (B) knockdown were measured by qPCR. (C-E) IFN-β promoter activity was assessed by luciferase assay following eIF2α (C), IFI16 (D) or STING (E) knockdown. (F) Schematic of hydrodynamic injection (HDI) of HBV plasmid and intraperitoneal Spiperone treatment in IFNAR-KO mice (G) The level of HBsAg, HBeAg, and HBV DNA in mouse serum samples were measured by ELISA or qPCR. (H) Hepatic pgRNA and cccDNA levels were measured using qPCR. (I and J) IHC images showing HBsAg expression (I) and confocal images showing HBcAg expression (Alexa Fluor 594) (J) in mouse liver tissue. The representative images are shown. (K) ChIP–qPCR analysis of cccDNA-associated histone acetylation in HepG2-NTCP-C4 cells. The significance of differences is indicated as *p < 0.05, **p < 0.01, and ***p < 0.001. ChIP, chromatin immunoprecipitation; HBV, Hepatitis B virus; IHC, Immunohistochemical.

To further evaluate the role of IFN-I signaling in the antiviral activity of Spiperone, we used an IFNAR-KO mouse model. Wild-type (WT) and IFNAR-KO mice were injected with 1.2x HBV plasmid via hydrodynamic injection (HDI) and the mice were subsequently treated with Spiperone or PBS for two weeks (Fig. 5F). In hydrodynamically injected C57BL/6 WT mice, Spiperone administration significantly reduced HBsAg, HBeAg, serum HBV DNA, hepatic pgRNA and cccDNA levels, whereas these antiviral effects were largely absent in IFNAR knockout mice (Fig. 5G, H). The results of immunohistochemical staining for HBsAg and immunofluorescence staining for HBcAg further supported these findings, showing clear reductions in antigen levels in WT livers but not IFNAR deficient livers after Spiperone treatment (Fig. 5I, J).

Taken together, these findings indicate that Spiperone suppresses HBV replication and cccDNA through PERK–eIF2α–ATF4–mediated ER stress, IFI16–STING–dependent DNA sensing, and downstream IFN-I signaling.

### 2.6 Spiperone modulates epigenetic modifications on HBV cccDNA minichromosomes

Since Spiperone reduced cccDNA levels through IFN-I signaling, we assessed whether it also altered the transcriptional state of cccDNA minichromosomes in HBV-infected HepG2-NTCP-C4 cells. As cccDNA transcription is tightly regulated by histone modifications ^46^, we performed ChIP–qPCR analysis to measure the levels of transcriptionally active histone marks associated with cccDNA. Spiperone treatment significantly reduced H4K5ac, H3K27ac, AcH3, and AcH4 markers of transcriptionally active chromatin (Fig. 5K). The reduction in H3K27ac (*p* < 0.001) and AcH4 (*p* < 0.001) was particularly pronounced, indicating that Spiperone shifts the chromatin environment of cccDNA toward a less transcriptionally permissive state. Together with the IFN-I-dependent effects of Spiperone on cccDNA reduction, these epigenetic changes further show that Spiperone limits cccDNA not only by decreasing its abundance but also by repressing its transcriptional output.

### 2.7 Spiperone reduces HBV replication and cccDNA persistence in an AAV-HBV1.04x mouse model

To assess the ability of Spiperone to reduce cccDNA levels *in vivo*, the AAV-HBV1.04× model was selected due to its ability to support de novo cccDNA formation in murine hepatocytes. Unlike integrated HBV transgenes, HBsAg production in this model occurs only from newly formed cccDNA^47^, allowing direct assessment of cccDNA-dependent viral activity. Male C57BL/6 mice were injected with AAV-HBV1.04x (8X 10^10^vg per mouse, *n* = 10 mice). Measurement of serum samples by ELISA at one-week post-injection confirmed robust HBsAg production, confirming the successful establishment of the cccDNA mouse model (Supplementary Fig. S14) (*p* < 0.001 vs Naive). One week later, the mice received an intraperitoneal injection of Spiperone (1.5 mg per kg, *n* = 5 mice) daily for 4 weeks (Fig. 6A). Spiperone treatment reduced serum HBsAg and HBeAg levels compared with those in AAV-HBV controls (Fig. 6B, C), indicating the suppression of viral antigen production. Serum HBV DNA levels were lower in the Spiperone group (Fig. 6D), and the hepatic levels of pgRNA were also decreased (Fig. 6E). Immunofluorescence detection of HBsAg in liver sections confirmed reduced antigen staining in the Spiperone-treated group, and immunohistochemical detection of HBcAg revealed a decreased number of HBV-positive hepatocytes (Fig. 6F, G). Quantitative PCR analysis of mouse liver cccDNA demonstrated a significant decline in cccDNA levels in Spiperone-treated mice (Fig.[6H) (*p* < 0.05). Consistently, southern blot analysis further confirmed the reduction in cccDNA levels (Fig.[6I), and moreover, cccDNA selective PCR assays showed fewer amplification products in Spiperone-treated livers without affecting non[selective amplification (Fig.[6J). Importantly, these results demonstrate that Spiperone reduces intrahepatic cccDNA levels *in vivo*, extending beyond its in vitro effects on HBV antigenemia and viral replication.

**Fig. 6.**
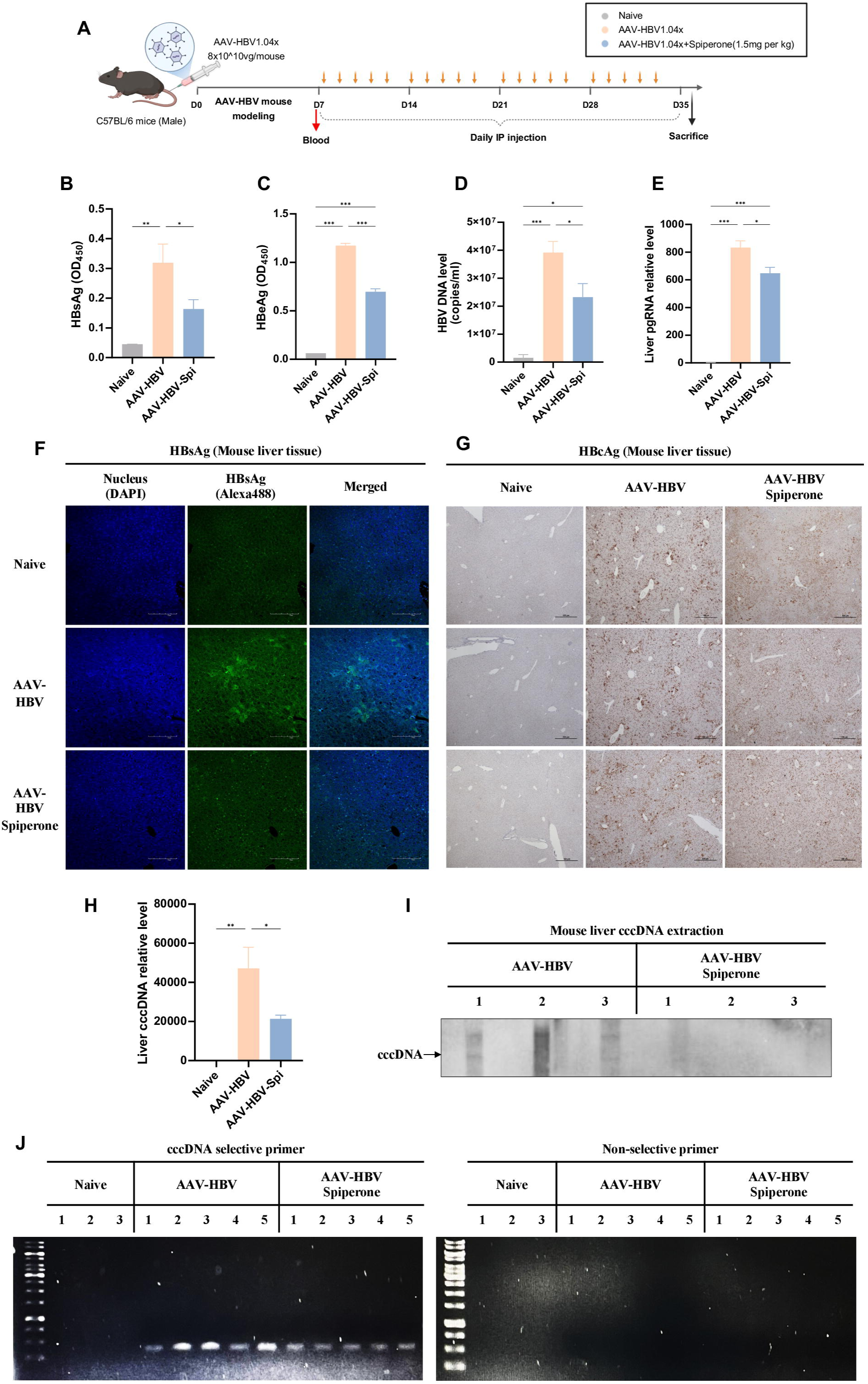
Spiperone reduces HBV replication and cccDNA persistence in an AAV-HBV1.04x mouse model. (A) Schematic illustration of intraperitoneal Spiperone administration in male AAV-HBV1.04× mice (male; *n* = 5 per each group). (B and C) The relative levels of HBsAg (B) and HBeAg (C) in mouse serum samples were measured by ELISA. (D and E) The levels of HBV viral DNA (D) in mouse serum and pgRNA (E) in mouse liver tissue were measured by qPCR. (F and G) Confocal images showing HBsAg expression (Alexa Fluor 488) (F) and IHC images showing HBcAg expression (G) in mouse liver tissue. The representative images are shown. (H) Quantification of cccDNA level was performed in mouse liver tissue by qPCR. (I) Southern blot analysis of liver tissue from AAV-HBV1.04x mice. (*n=3* per each group) (J) The cccDNA level was assessed by agarose gel electrophoresis using Hirt-extracted DNA amplified with cccDNA selective or non-selective primers. The significance of differences is indicated as *p < 0.05, **p < 0.01, and ***p < 0.001. HBV, Hepatitis B virus; IHC, Immunohistochemical.

In the AAV-HBV1.04× mouse model, where viral protein expression depends on de novo cccDNA formation, Spiperone reduced HBV replication and decreased cccDNA levels. These results suggest that Spiperone targets key steps in the HBV life cycle and may represent a promising therapeutic candidate for chronic HBV infection.

## 3. Discussion

In this study, we identify Spiperone as a host-directed antiviral agent that suppresses HBV replication and reduces cccDNA through coordinated activation of ER stress–linked innate immune signaling and epigenetic regulation. Mechanistically, Spiperone activates the PERK–eIF2α–ATF4 axis, promotes mitochondrial stress and cytosolic release of oxidized mtDNA, and subsequently engages IFI16–STING–dependent IFN-I signaling. This cascade culminates in suppression of viral transcription and reduced enrichment of activating histone marks on the cccDNA minichromosome, indicating both quantitative reduction and functional silencing of cccDNA.

These findings provide mechanistic insight into how host stress responses can be leveraged to control HBV persistence. While ER stress has been implicated in antiviral signaling, its functional linkage to cccDNA regulation has remained incompletely defined. Our data support a model in which ER stress–induced mitochondrial perturbation generates immunostimulatory mtDNA that is sensed by IFI16, thereby coupling cellular stress to innate immune activation. The resulting IFN-I response not only restricts viral intermediates but is also associated with epigenetic remodeling of cccDNA, shifting the chromatin state toward transcriptional repression. This integration of stress signaling, DNA sensing, and chromatin regulation represents a distinct host-mediated mechanism for controlling cccDNA activity.

Within the current landscape of cccDNA-targeting strategies, several features distinguish Spiperone from existing modalities. Gene-editing approaches, including CRISPR-based systems, can directly target cccDNA but face significant translational challenges, such as efficient delivery to hepatocytes, incomplete editing, and potential off-target or genomic rearrangement risks. Capsid assembly modulators primarily inhibit nucleocapsid recycling and de novo cccDNA formation, with limited impact on the pre-existing cccDNA pool ^48^. In contrast, Spiperone operates through activation of endogenous host pathways that converge on innate immune signaling and chromatin remodeling, thereby suppressing both viral replication and the transcriptional competence of established cccDNA. This distinction highlights the potential of host-directed strategies to complement existing antiviral classes by targeting regulatory layers of cccDNA persistence ^49^.

Importantly, the antiviral activity of Spiperone was consistently observed across multiple experimental systems, including stable replication models, bona fide infection systems, and in vivo mouse models supporting de novo cccDNA formation. The reduction of viral markers following both intraperitoneal and oral administration underscores its pharmacological robustness and translational feasibility. In addition, the observed synergy with tenofovir suggests that Spiperone may complement nucleos(t)ide analogs by targeting cccDNA through mechanisms distinct from inhibition of reverse transcription, supporting its potential use in combination regimens aimed at more effective reservoir control.

From a translational perspective, drug repurposing offers a practical advantage, as Spiperone has an established safety and pharmacokinetic profile. The dosing used in our study falls within ranges previously shown to be tolerated in rodents and may be clinically relevant based on allometric scaling ^50–52^. These features support further development of Spiperone, or agents targeting similar pathways, for therapeutic application in chronic HBV infection.

Several limitations should be considered. First, the precise upstream molecular target through which Spiperone initiates ER stress remains to be defined. Second, validation in primary human hepatocytes and long-term models will be necessary to assess durability and safety. Finally, the therapeutic potential of Spiperone in combination with other antiviral or immunomodulatory agents warrants systematic investigation.

In conclusion, Spiperone suppresses HBV replication and cccDNA persistence through a host-directed mechanism integrating ER stress, innate immune activation, and epigenetic regulation. These findings provide a conceptual framework for targeting cccDNA via stress-induced immune pathways and support the development of host-targeted combination strategies toward a functional cure for chronic HBV infection.

## 4. Materials and Methods

### 4.1 Cell cultures and Transfection

HepG2.2.15 cells (#SCC249) were purchased from Sigma –Aldrich (St. Louis, MO, USA) and maintained in Dulbecco’s Modified Eagle Medium (DMEM) supplemented with 10% fetal bovine serum (FBS) and Penicillin-streptomycin (PS). HepG2-NTCP-C4 cells 33 (kindly gifted by Wakita et al.) were cultured in DMEM/F12 medium supplemented with 10% fetal bovine serum (FBS), 100U/ml Penicillin-streptomycin (PS), 10 mM HEPES, and 5 µg/ml insulin solution. HepaRG cells (HPR116) were obtained from Biopredic International (Saint-Gregoire, France). The cells were thawed using thawing/plating medium (ADD670C; Biopredic, Saint-Gregoire, France) and subsequently cultured in basal hepatic cell medium (MIL600C; Biopredic) with additives for maintenance/metabolism HepaRG medium (ADD620C; Biopredic). For the reporter assay, hMH55-293-ISRE cells containing integrated interferon-sensitive response element (ISRE) genes with a luciferase reporter tag on chromosomes were utilized.

### 4.2 Mouse Experiments

Six- to eight-week-old male HBV transgenic (TG) mice (IACUC number SNU-250310-5), IFN-I receptor α-chain knock-out (IFNAR KO) mice (IACUC number SNU-220401-3), and C57BL/6 mice (IACUC number SNU-241202-1-1) were used in this study. HBV Transgenic mice (generated by Macrogen, Inc. IFNAR KO 129/SvEv mice ^53^) were kindly provided by Dr. Heung Kyu Lee (Korea Advanced Institute of Science and Technology) and backcrossed with C57BL/6J more than ten generations. To establish HBV cccDNA model, C57BL/6 mice were injected via the tail vein with recombinant Adeno-associated virus (AAV)-HBV1.04 (8×10^10^vg diluted in 200 μL Phosphate-buffered saline (PBS)). The recombinant AAV-HBV1.04 plasmids were kindly provided by Dr. Yuchen Xia ^47^. All experiments utilizing animals in this study were approved by the Seoul National University Institutional Animal Care and Use Committee. For HBV infection, IFNAR KO mice or C57BL/6 mice were hydrodynamically injected via the tail vein with 10 μg of pAAV-HBV plasmid carrying the full-length HBV genotype C genome, an amount equivalent to 10% of the mouse body weight. The total volume was delivered within 5–8 s. All the mice used in this study were anesthetized in an induction chamber with isoflurane (3% in oxygen). All animal experiments were approved by the Institutional Animal Care and Use Committee of Seoul National University.

### 4.3 Drug treatment

For the in vivo assay, various mouse models were used in this study. HBV transgenic mice were administered Tenofovir disoproxil fumarate (TDF) (Sigma–Aldrich, SML1794) (500μg/mouse) or Spiperone (MedchemExpress, Monmouth Junction, NJ, USA, HY-B1371) (1.5mg/kg) daily for eight weeks via either intraperitoneal injection (IP) or oral administration. IFNAR KO mice or C57BL/6 mice via hydrodynamic injection (HDI) were administered PBS or 1.5mg/kg Spiperone daily for two weeks. For the cccDNA mouse model, the mice received 1.5mg/kg Spiperone daily for four weeks. For the in vitro assay, HepG2.2.15 cells, as well as HepG2 cells transfected with the pHBV-1.2x-wild-type genotype C plasmid were treated with Tenofovir (TDF) (5 μM) or Spiperone (10 μM) to assess the antiviral effect.

### 4.4 HBsAg and HBeAg ELISA assay

Supernatants from cells and serum samples from mice were collected. Blood samples were obtained at the indicated time points, and sera were prepared by centrifugation at 13000 rpm for 15 min at 4°C. Hepatitis B surface antigen (HBsAg) and hepatitis B e antigen (HBeAg) levels in the supernatants were quantified using the Hepatitis B Virus HBsAg ELISA Kit (Abnova, Taipei, Taiwan) and the HBeAg ELISA kit (Novus Biologicals, Centennial, CO, USA), respectively, according to the manufacturer’s instructions.

### 4.5 Quantitative PCR (qPCR) for HBV DNA quantification

HBV DNA was extracted using the QIAamp DNA Blood Mini Kit (QIAGEN, Hilden, Germany) following the manufacturer’s instructions and was quantified by quantitative PCR (qPCR) using a SensiFAST™ SYBR® Lo-ROX Kit (Bioline, London, UK). The sequences of primers used for the amplification are shown in Table S1.

### 4.6 HBV pgRNA Extraction and Quantification

Total RNA was extracted from cell pellets or mouse liver tissue using TRIzol reagent (Invitrogen, Carlsbad, CA, USA) according to the manufacturer’s instructions. Cell pellets or 50 mg of liver tissue were fully homogenized, and precipitated using chloroform/isopropanol method. RNA samples were treated with RQ1 DNase (Promega, Southampton, UK) for 60 min at 37°C prior to the addition of 1 μl of stop solution and incubation at 65 °C for 10 min for DNase inactivation. The RNA was stored at −80°C until further use. For analysis of viral pgRNA, 1 μg of total RNA was reverse-transcribed and amplified using a SensiFAST™ cDNA Synthesis Kit (Bioline). Subsequently, 1μl of cDNA was quantified via RT-qPCR, with glyceraldehyde 3-phosphate dehydrogenase (GAPDH) serving as the internal control for normalization. The sequences of primers are listed in Table S1.

### 4.7 HBV cccDNA extraction and Quantification

HepG2.2.15 cells treated with TDF or Spiperone were lysed in buffer containing NP-40 at 4°C for 10 min. After centrifugation, the nuclear pellet was resuspended in buffer containing SDS, sonicated, and digested with proteinase K overnight. DNA was extracted by phenol-chloroform method, precipitated with ethanol, and treated with Plasmid Safe DNase I (PSAD) for 45 min. The reaction was terminated by heating at 70°C for 30 min. cccDNA levels were quantified by qPCR, with mitochondrial DNA (mtDNA) for normalization. For analysis of intrahepatic cccDNA, 50 mg of liver tissue was fully homogenized in lysis buffer (75 mM NaCl, 25 mM EDTA) and incubated at 55 °C after mixing with 10% SDS and Proteinase K solution. The homogenate was extracted twice with an equal volume of Tris-saturated phenol/chloroform, and the supernatant was precipitated with 3M Sodium acetate (NaAC) (pH 5.2) and isopropanol. The pellet was subsequently washed with 70% ethanol and dissolved in Tris-EDTA (TE) buffer. DNA was treated PSAD and the reaction was terminated by heating at 70°C for 30 min as described above. cccDNA levels were quantified by qPCR, with mtDNA used as the internal control. The sequences of primers are shown in Table S1. To detect HBV cccDNA on electrophoresis, DNA samples were amplified using cccDNA selective primers and non-cccDNA-selective primers and were then separated on an agarose gel. ^54^

### 4.8 mRNA level of ER stress-related and IFN-I-related genes

Total RNA was extracted from cell pellets using TRIzol reagent (Invitrogen) according to the manufacturer’s instructions. The RNA samples were treated with RQ1 DNase (Promega) for 60 min at 37°C, followed by the addition of 1 μl of stop solution and incubation at 65°C for 10 min for DNase inactivation. The RNA was stored at −80°C until further use. To measure the transcript levels of endoplasmic reticulum (ER) stress-related and IFN-I-related genes, 1μg of total RNA was reverse-transcribed and amplified using a SensiFAST™ cDNA Synthesis Kit (Bioline) Subsequently, 1μl of cDNA was quantified via RT-qPCR with specific primer sets (Table S1), with GAPDH for RNA normalization. IFN-I-related genes are obtained from AccuPower® qPCR Array System: Immune qPCR Panel Kit (Bioneer, Daejeon, South Korea)

### 4.9 Western blot (WB) assay

Proteins were extracted from HepG2.2.15 cells using Radioimmunoprecipitation assay (RIPA) buffer (Invitrogen) supplemented with protease inhibitors (cOmplete^™^ Mini EDTA-free Protease Inhibitor Cocktail) (Roche, Basel, Switzerland) and phosphatase inhibitors (PhosSTOP^™^) (Roche) and quantified via a standard Bicinchoninic acid assay (BCA). Equal amounts of protein (10 µg) were separated by sodium dodecyl sulfate-polyacrylamide gel electrophoresis (SDS-PAGE) and transferred onto nitrocellulose membranes. The membranes were blocked with 5% bovine serum albumin (BSA) in Tris-Buffered Saline with tween (TBS-T) and incubated overnight at 4°C with primary antibodies (diluted 1/1000). After being washed, the membranes were incubated with horseradish peroxidase (HRP)-conjugated secondary antibodies (diluted 1/10000) for 2 h at room temperature. Protein bands were visualized using enhanced chemiluminescence (ECL) and imaged with a ChemiDoc system, and band densities were then quantified using ImageJ software. The list of antibodies used in this study was shown in Table S2.

### 4.10 Southern Blot for cccDNA detection

DIG DNA Labeling and Detection Kit (Roche) was used to detect the cccDNA in liver tissue and cell pellets. cccDNA was extracted as described above. cccDNA was amplified using phi29 DNA polymerase (Thermo Scientific, Carlsbad, CA, USA). Briefly, the extracted DNA was amplified with phi29 DNA polymerase, Exo-resistant random primers (Thermo Scientific) and dNTP mix for 4 h at 30°C followed by 65°C for 10 min. After amplification, PCR was performed using the P1-P2 primer set (Table S1), which targets the full-length HBV genome. This protocol was adapted from the method of Dr. Lebossé, Fanny ^55^. Gel running was performed on 1.2% agarose gel with 50 V, and subsequently transferred to a Hybond N+ nylon membrane (Amersham, Little Chalfont, UK). The membrane was hybridized overnight at 42°C-45°C with a 1.2X wild-type HBV full-genome probe digested with XhoI. After hybridization, the membrane was washed with washing buffer, incubated with antibody solution, and visualized using detection buffer and color substrates provided in the kit.

### 4.11 Immunofluorescence (IF) and Confocal Microscopy

Mouse liver tissues were fixed, paraffin-embedded, and sliced into 4 μm-thick sections prior to deparaffinization. HepG2.2.15 cells were cultivated into 4-chamber glass slides (Nunc, Roskilde, Denmark), treated with TDF or Spiperone and fixed with methanol. Liver tissue sections and cultured cells were incubated with anti-HBcAg (Abcam, ab8637, 1/200), anti-HBsAg (Santa Cruz, sc-53299, 1/200), anti-ATF4 (Santa Cruz, sc-390063, 1/200), anti-8-OHdG (Santa Cruz, sc-393870, 1/200) overnight at 4°C and then with Alexa 488 – or 594-conjugated secondary antibodies (diluted 1/1000) for 1 h at room temperature in Phosphate-buffered saline with tween (PBS-T) containing 1% BSA. Nuclear staining was performed using 4′,6-diamidino-2-phenylindole (DAPI)-containing mounting medium (VECTASHIELD, H-1200; Vector Laboratories, Inc., Burlingame, CA, USA). The list of antibodies used in this study was shown in Table S2.

### 4.12 Immunohistochemistry (IHC)

Liver tissues were fixed, paraffin-embedded, and sliced into 4 μm-thick sections prior to deparaffinization. The sections were incubated in xylene and rehydrated through a graded series of ethanol concentrations. Antigen retrieval was performed by heating the slides in sodium citrate buffer (pH 6.0) at 95°C for 20 min, followed by cooling at room temperature. Endogenous peroxidase activity was blocked by incubating the sections with 3% hydrogen peroxide for 10 min. After being washed with 0.025% Triton X-100 in PBS, the sections were incubated overnight at 4°C with anti-HBcAg (Sigma–Aldrich, 216A-15, 1/200), anti-HBsAg (Santa Cruz, sc-53299, 1/200), anti-p-PERK (Santa Cruz, sc-32577, 1/200), anti-ATF4 (Santa Cruz, sc-390063, 1/200) diluted in PBS containing 10% FBS and 1% BSA. The next day, the sections were incubated with species-specific HRP-conjugated secondary antibodies (diluted 1/1000) for 1 h at room temperature. Staining was visualized using the chromogenic substrate 3,3′-diaminobenzidine (DAB), and the reaction was monitored under a microscope to prevent overdevelopment. Finally, the slides were counterstained with hematoxylin, dehydrated and mounted for examination under a light microscope.

### 4.13 Chromatin immunoprecipitation (ChIP)-qPCR assay

To analyze histone post-translational modifications (PTMs) on cccDNA-associated chromatin, a SimpleChIP Enzymatic Chromatin IP Kit (Cell Signaling Technology, Danvers, MA, USA, 9003) was used. HepG2-NTCP-C4 cells were infected with HBV WT particles. The cell pellets were subsequently washed with PBS, crosslinking was performed with formaldehyde, and the reaction was quenched with glycine. After cross-linking, the cells were lysed and treated with nuclease, and the nuclear pellets were sonicated for chromatin digestion. Protein-DNA complexes were immunoprecipitated using magnetic beads and antibodies against Histone H4 lysine 5 acetylation (H4K5ac), Histone H3 lysine 27 acetylation (H3K27ac), Acetylated Histone H3 (AcH3) or Acetylated Histone H4 (AcH4), or with normal IgG as a negative control. The beads were subsequently washed with wash buffer to reduce nonspecific binding, eluted with elution buffer, heated for 30 min, and treated with proteinase K, after which the DNA was purified using the DNA wash buffer provided in the kit. A 2% input DNA control was processed in parallel. The purified DNA was analyzed by qPCR using the cccDNA primer listed in Table S1, and the values were normalized to input DNA.

### 4.14 Statistical Analysis

All the statistical analyses were performed using GraphPad Prism software (version 9.0; GraphPad Software, San Diego, CA, USA). Statistical analyses were performed using one-way or two-way ANOVA. Post-hoc analysis was performed using Tukey’s multiple-comparison test. The p value of significance was designated at p < 0.05 (*), 0.01 (**), or 0.001 (***). The data are presented as the mean ± S.D. of three independently performed experiments.

A detailed description of the materials and methods is provided in the supplementary data.

## Acknowledgments

This work was supported by the Korea Health Technology R&D Project through the Korea Health Industry Development Institute (KHIDI), funded by the Ministry of Health & Welfare, Republic of Korea (RS-2024-00456525), the Basic Science Research Program through the National Research Foundation of Korea (NRF) funded by the Ministry of Education (RS-2025-00553721), and a grant no 16-2023-0011 from Seoul National University Bundang Hospital (SNUBH) Research Fund. We thank Dr. Koichi Watashi (National Institute of Infectious Disease, Tokyo, Japan) for providing the HepG2-NTCP-C4 cells.

## Author contributions

Bum-Joon Kim contributed to the conception and design of the study. Junghwa Jang, Ziyun Kim, Eunseo Kim, Jisung Park, Yu-Min Choi performed the laboratory work and organized the database. Bum-Joon Kim, Junghwa Jang, Ziyun Kim, Yu-Min Choi wrote and revised the manuscript. All the authors read and approved the submitted version.

## Conflict of interest

The authors declare that they have no competing financial interests.

## Ethics approval and consent to participate

This animal study was approved by the Institutional Animal Care and Use Committee (IACUC) of Seoul National University (Approval No. SNU-241202-1-1).

## Availability of data and materials

All the data generated or analyzed during this study are included in this article and its additional files.

---

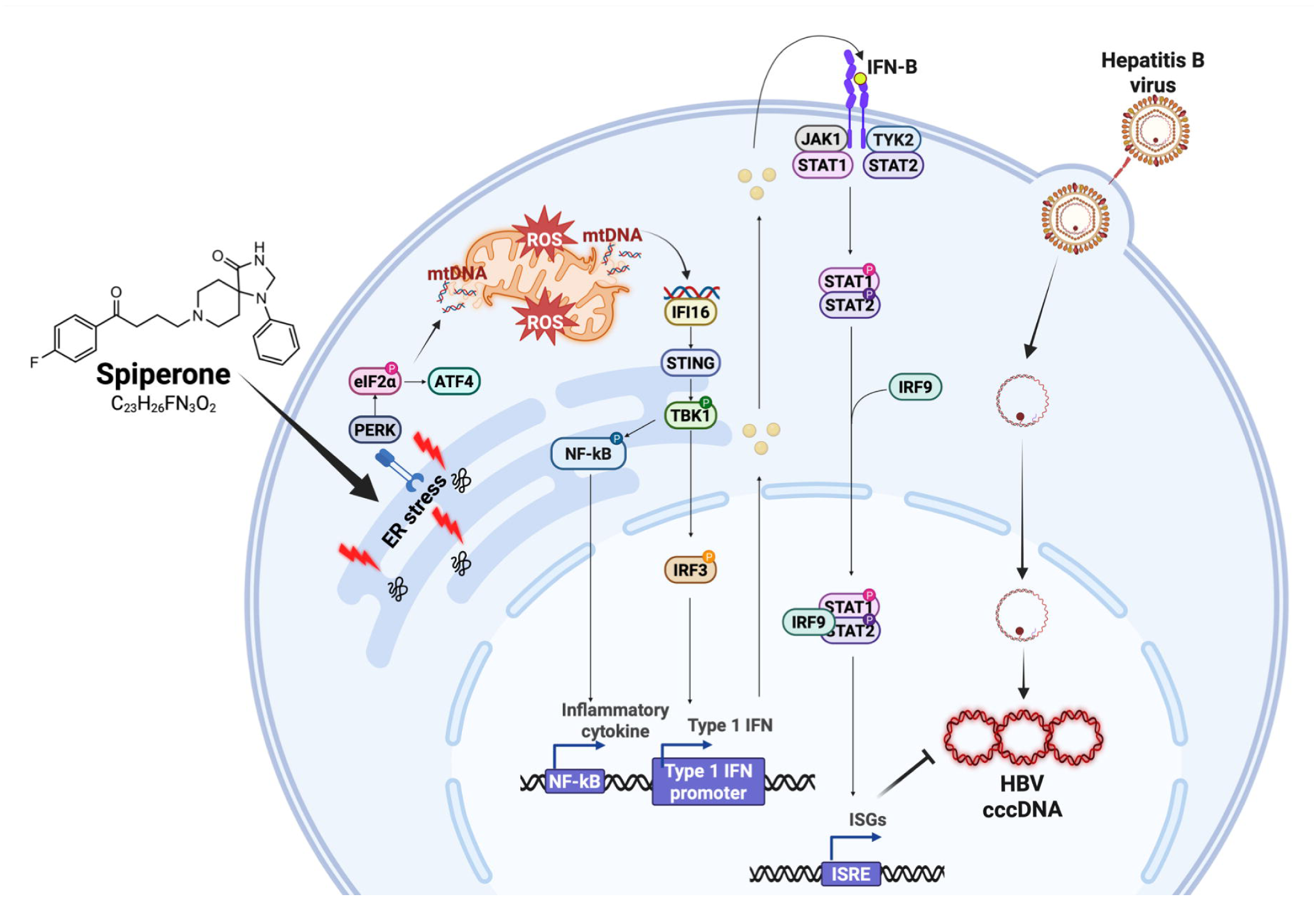

## Notes

### Competing Interest Statement

The authors have declared no competing interest.

