## Supplementary materials, methods, Figures and tables for "Spiperone targets HBV cccDNA via ER stress–induced innate immune activation and epigenetic silencing"

**Table of contents**

Supplementary materials and methods ………………………………………………… 2

Supplementary Figures ……………………………………………………………….... 5

Supplementary Tables ………………………………………………………………… 20

**Supporting Materials**

**Supplemental materials and methods**

1. **siRNA transfection**

For siRNA transfection, eukaryotic initiation factor 2 α (eIF2α)-siRNA (sc-78173; Santa Cruz Biotechnology, Dallas, TX, USA), interferon gamma-inducible protein 16 (IFI16)-siRNA (sc-35633; Santa Cruz Biotechnology) and Stimulator of interferon genes (STING)-siRNA (sc-92042; Santa Cruz Biotechnology) transfections were performed using Lipofectamine 3000 (Invitrogen) according to the manufacturer’s instructions. Scramble siRNA (sc-37007; Santa Cruz Biotechnology) was used as a control. After drug treatment or siRNA transfection, the supernatants and cell pellets were collected for Enzyme-linked immunosorbent assay (ELISA) and Reverse transcription quantitative PCR (RT-qPCR) analyses.

2. **The synergistic evaluation of drugs**

The synergistic effect was assessed by HBsAg levels in the sample using the Hepatitis B Virus HBsAg ELISA Kit (Alpha diagnostics Intl. Inc. Texas, USA and Abnova, Taiwan) using the ZIP model, and ZIP scores were calculated with the SynergyFinder algorithm version 3.0 (<https://synergyfinder.org/>)^1^. Consistent with previous report ^2^, a ZIP score greater than +5 was regarded as synergistic and adopted in this study.

3. **Cytosolic mtDNA measurement**

The release of mtDNA in the cytosol was measured as previously described ^3^. HepG2.2.15 cells were seeded into 6-well plate and treated with 5 μM TDF or 10 μM Spiperone. The cell pellets were lysed in a mild lysis buffer containing NP-40 and 1x protease inhibitor cocktail, then divided into two tubes. One tube was used for total DNA extraction, and the other tube was incubated on ice for 15 min. After incubation, the mixture was centrifuged at 13,500 xg for 15min at 4 °C. The supernatant was transferred to a new tube as the cytosolic fraction, which was used for cytosolic mtDNA extraction. DNA was extracted from both the cytosolic and total fractions using a AccuPrep® Genomic DNA Extraction Kit (Bioneer). We evaluated the mtDNA (ND1, ND2) level in the cytosolic fraction and 18S rRNA (housekeeping gene) level in the total DNA fraction. The release of mtDNA into the cytosol was quantified by qPCR. The relevant primer sequences are shown in Table S1.

4. **MitoSox Measurement**

Intracellular Reactive oxygen species (ROS) levels were measured using the the mitochondrial superoxide indicator MitoSOX™ Red (Invitrogen, M36008). HepG2.2.15 cells were seeded into 6-well plate and treated with 5 μM TDF or 10 μM Spiperone. At the indicated time, 1 μM mitochondria-targeted superoxide (MitoSOX) was added to the cell pellets, which were then incubated for 10-20 min. After incubation, the pellets were washed twice with pre-warmed PBS containing 1 mM EDTA and 10% FBS. Intracellular ROS was measured by flow cytometry, with rotenone used as a positive control.

5. **Type I IFN Reporter Assay**

The type I IFN level was measured using hMH55-293-Interferon-stimulated response element (ISRE) reporter cells, as previously described ^4^. Briefly, cell culture supernatants from TDF- or Spiperone-treated cells were incubated with hMH55-293-ISRE cells for 12–24 h. After incubation, the reporter cells were lysed using Reporter Lysis Buffer (Promega). The lysates were then mixed with Luciferase Assay Reagent (Promega), and luminescence was measured using a TECAN plate reader.

6. **RNA sequencing**

For RNA sequencing analysis, HepG2-NTCP-C4 cells were infected with WT HBV particles, and replaced with or without Spiperone 10μM in DMEM/F12 supplemented with 2% FBS, Penicillin-streptomycin (PS), HEPES, and insulin solution. After 12 h, total RNA was extracted from cell pellets using using TRIzol reagent (Invitrogen) according to the manufacturer’s instructions. RNA samples were analyzed by Ebiogen (Korea). Data acquisition and processing were performed using ExDEGA program provided by Ebiogen. Fold changes (PBS vs Spiperone) were calculated for each sample. A heatmap of differentially expressed genes (Fold change (FC) <0.5 or >2, p-value < 0.05) and pathway over-representation of significantly upregulated genes (p-value <0.05 and FC >2) were generated by ExDEGA program and with the InnateDB web resource.

**Supplemental Figure Legends**

**(A)	(B)**

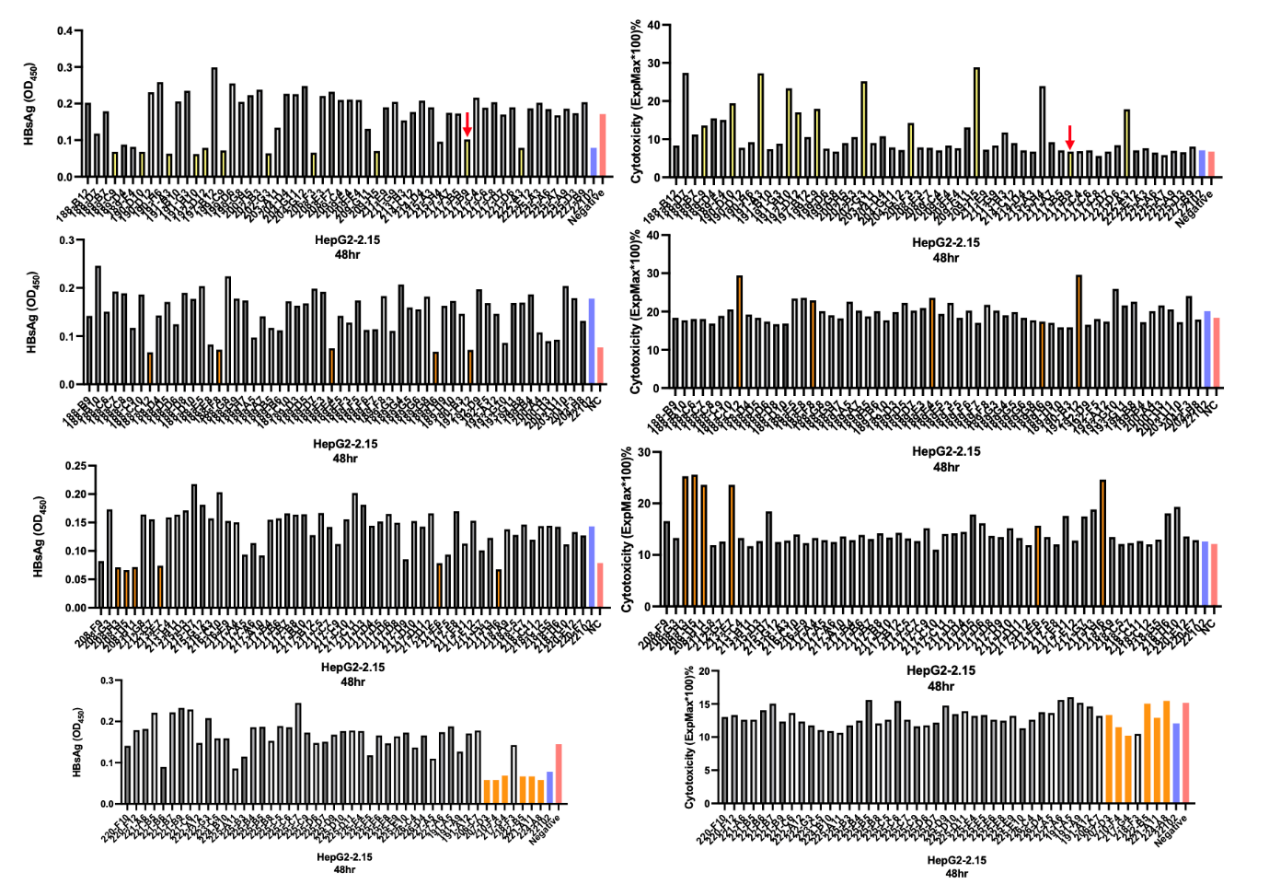

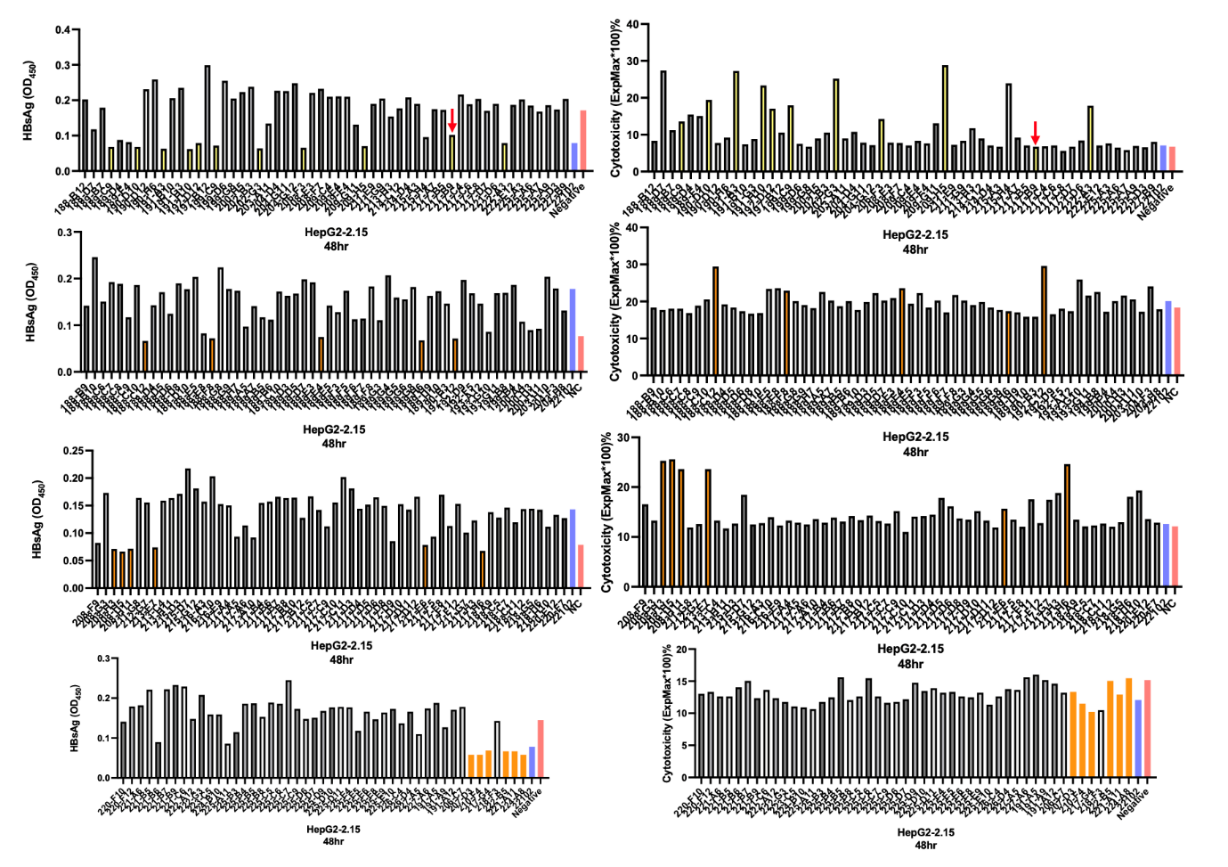

**Supplementary Figure 1. Primary screening for HBV cccDNA inhibitor candidates**.

A total of 500 compounds preselected from 3,200 clinical-stage and approved drugs obtained from the Korea Chemical Bank on the basis of cytotoxicity were tested in HepG2.2.15 cells at 10 μM for 48 h. (A) HBsAg levels in the supernatants were measured by ELISA. (B) Cell cytotoxicity was assessed using an LDH release assay (CytoTox 96® Non-Radioactive Cytotoxicity Assay, Promega) performed on culture supernatants collected at 48 h. HBV, Hepatitis B virus; HBsAg, HBV surface antigen; LDH, Lactate Dehydrogenase; cccDNA, covalently closed circular DNA.

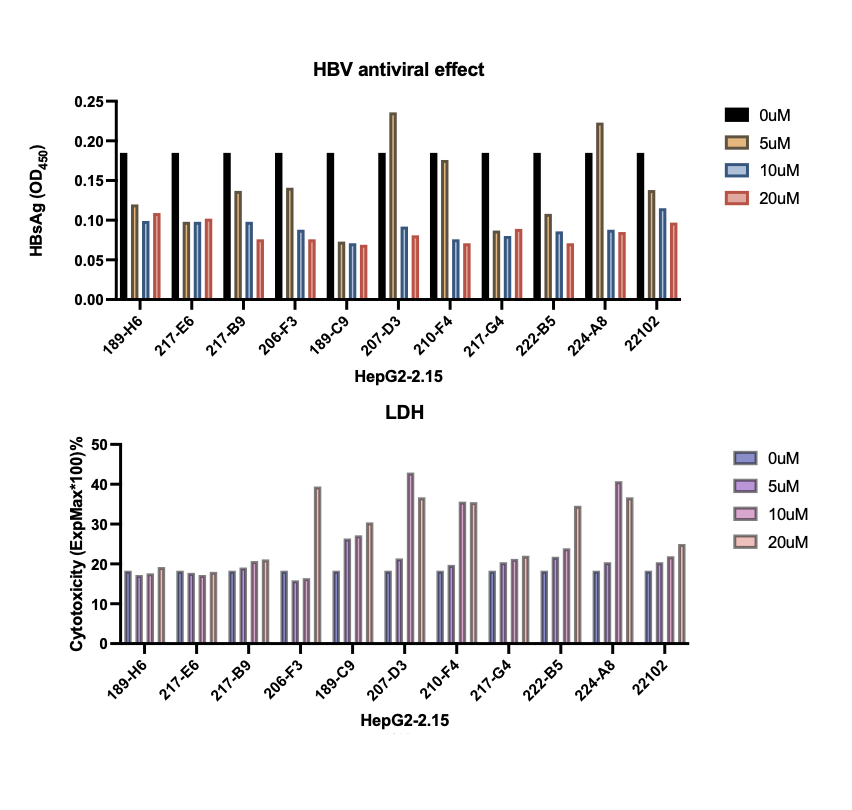
**(A)**

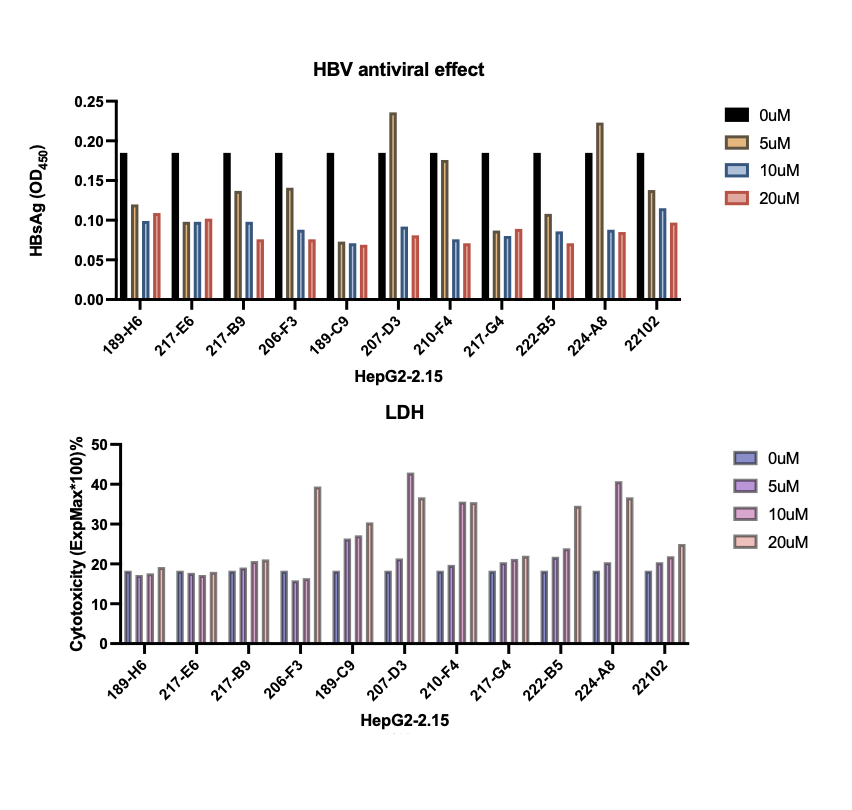
**(B)**

**Supplementary Figure 2. Dose-dependent effects of selected compounds on HBV replication and cytotoxicity**

Ten compounds identified via the primary screening were tested in HepG2.2.15 cells at 5, 10, and 20 μM for 48 h. (A) HBsAg levels in culture supernatants were measured by ELISA. (B) Cytotoxicity was assessed by an LDH release assay (Promega). HBV, Hepatitis B virus; HBsAg, HBV surface antigen; LDH, Lactate Dehydrogenase

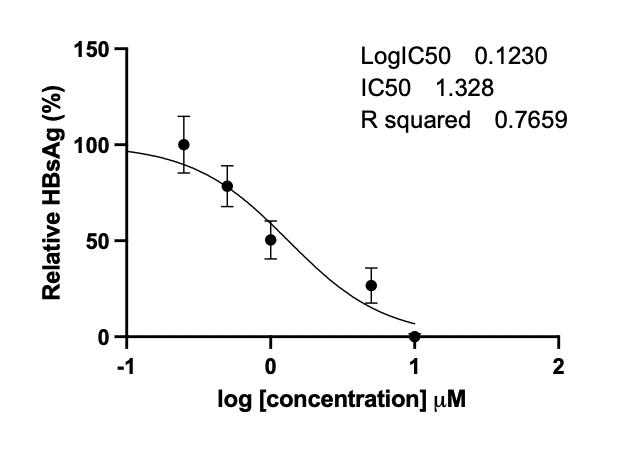

**Supplementary Figure 3. Dose-dependent suppression of HBsAg secretion in HepG2.2.15 cells**

HepG2.2.15 cells were treated with the indicated concentrations of Spiperone (0.25, 0.5, 1, 5, 10 μM) for 48 h. HBsAg levels in the supernatants were quantified using a Hepatitis B Virus HBsAg ELISA Kit (Abnova), and the absorbance was measured at OD 450nm in a TECAN plate reader. HBsAg, HBV surface antigen.

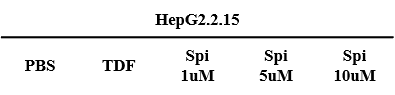

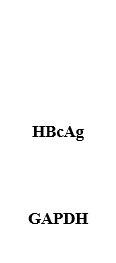

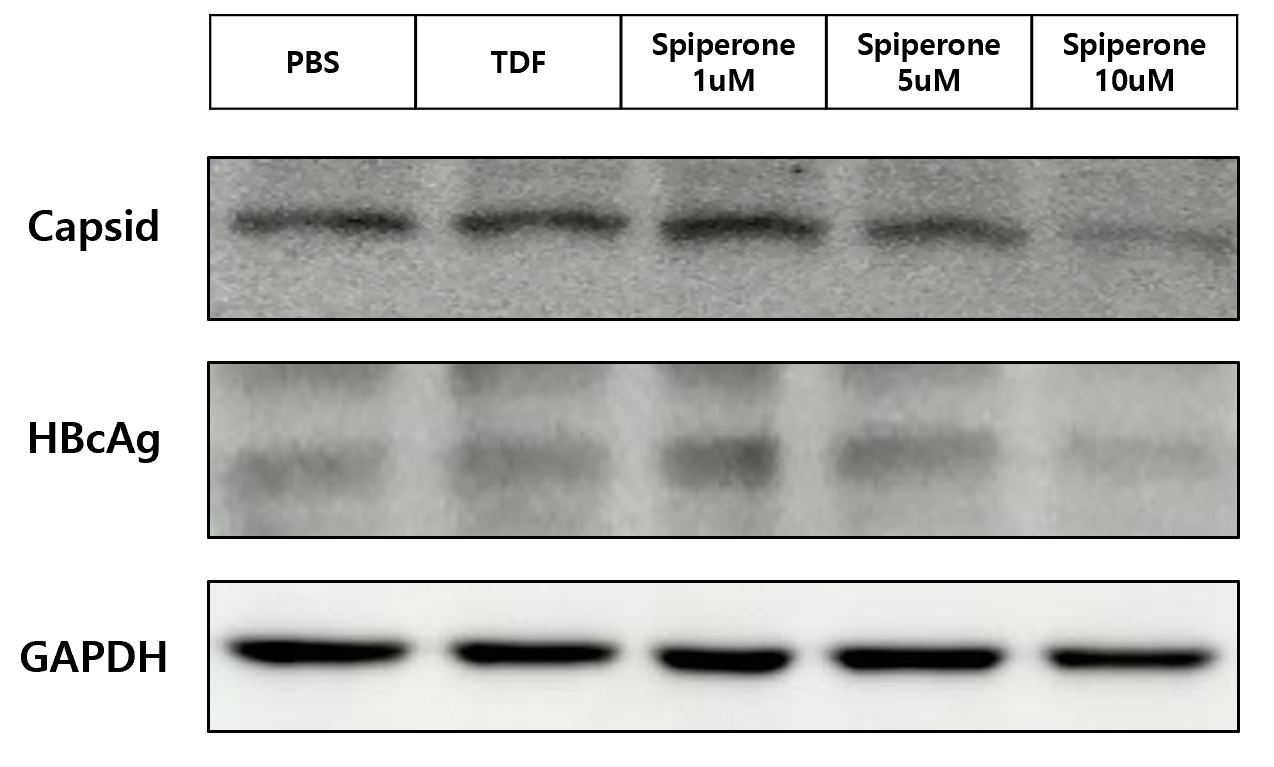

**Supplementary Figure 4. Capsid expression in response to Spiperone treatment.**

HepG2.2.15 cells were treated with PBS, TDF or Spiperone (1, 5, 10 μM) for 48h. HBcAg expression was detected by western blotting using an anti-HBc antibody. TDF, tenofovir disoproxil fumarate; HBcAg, HBV core antigen.

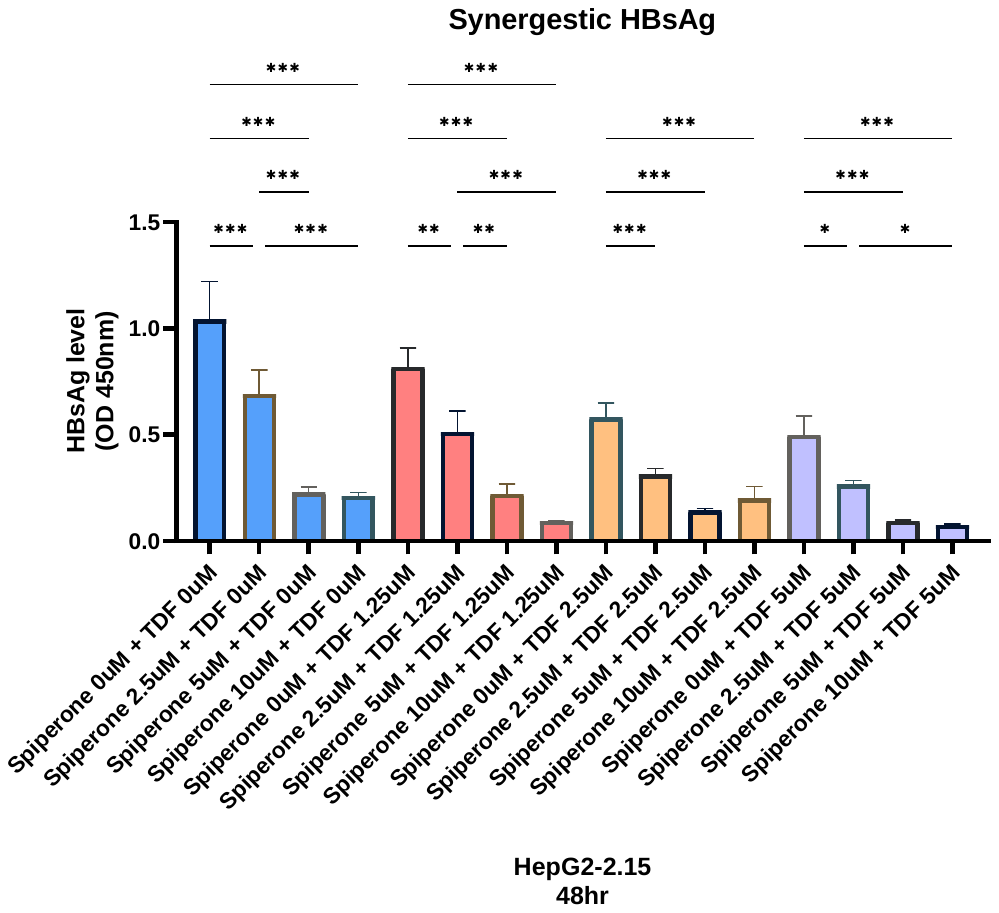

**TDF 1.25 μM**

**TDF 0 μM**

**TDF 2.5 μM**

**TDF 5 μM**

**Supplementary Figure 5. The synergistic effect of TDF and Spiperone**

HepG2.2.15 cells were treated with increasing concentrations of TDF (0-5 μM) or Spiperone (0-10 μM) for 48 hours. HBsAg expression in the supernatants were determined by ELISA. Data indicate the mean ± S.D. of three independently performed experiments. * *p* < 0.05, ** *p* < 0.01 and *** *p* < 0.001. TDF, tenofovir disoproxil fumarate; HBsAg, HBV surface antigen.

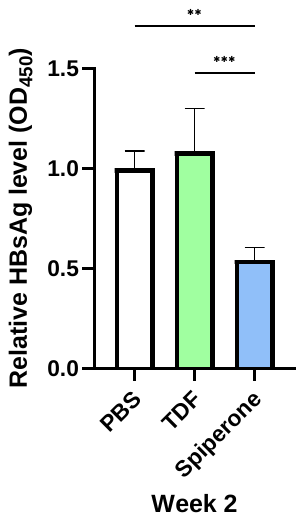

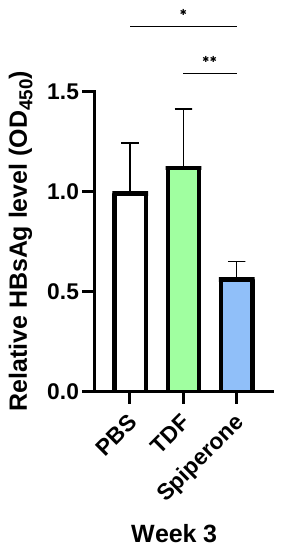

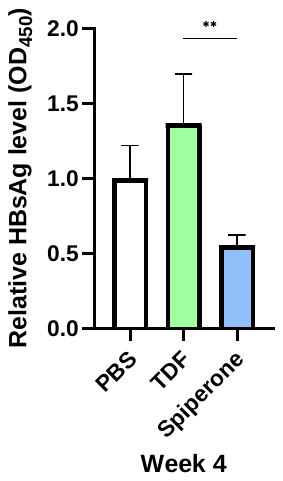

**Supplementary Figure 6. HBsAg levels in HBV TG mice treated with PBS, TDF, or Spiperone**

HBV TG mice (male) were administered PBS, TDF (5 mg per mouse) or Spiperone (1.5 mg per kg) daily by intraperitoneal (IP) injection. HBsAg expression in mouse serum samples were measured by ELISA at weeks 2, 3, and 4. Data indicate the mean ± S.D. of three independently performed experiments. * *p* < 0.05, ** *p* < 0.01 and *** *p* < 0.001. IP, Intraperitoneal injection; HBsAg, HBV surface antigen.

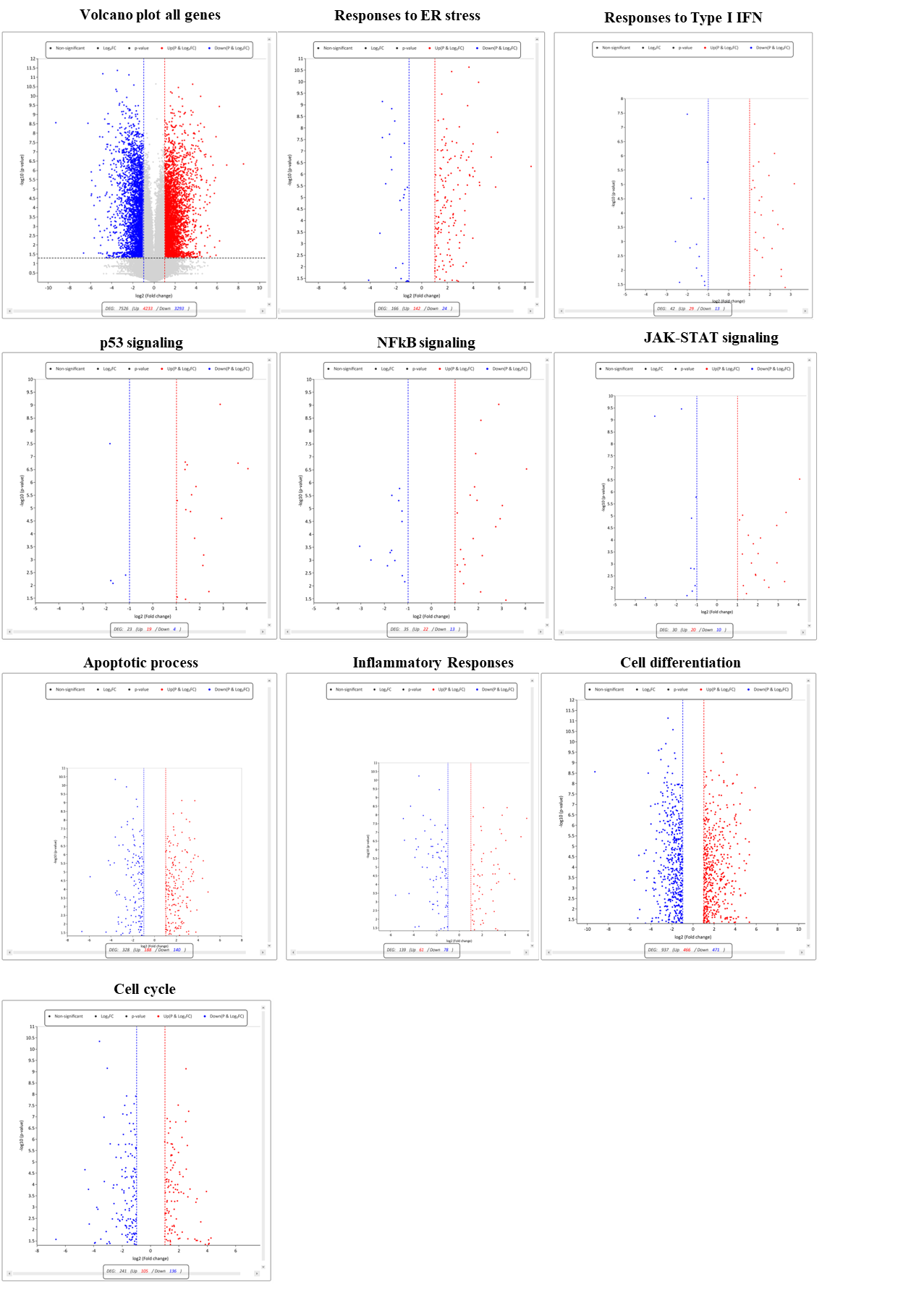

**Supplementary Figure 7. Volcano plots of the RNA-seq data for HepG2.2.15 cells following Spiperone treatment.**

Differentially expressed genes (DEGs) were identified by RNA sequencing after treatment with Spiperone. The red and blue dots indicate significantly upregulated and downregulated genes, respectively, based on cut-off values of |log2-fold change| ≥ 1 and adjusted P value < 0.05. Each panel corresponds to comparisons between Spiperone-treated and PBS-treated cells under the indicated condition**.**

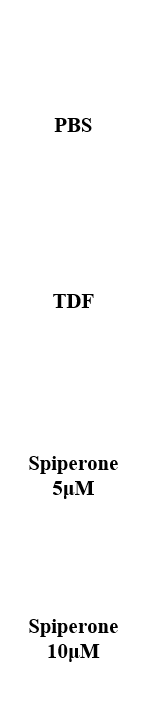

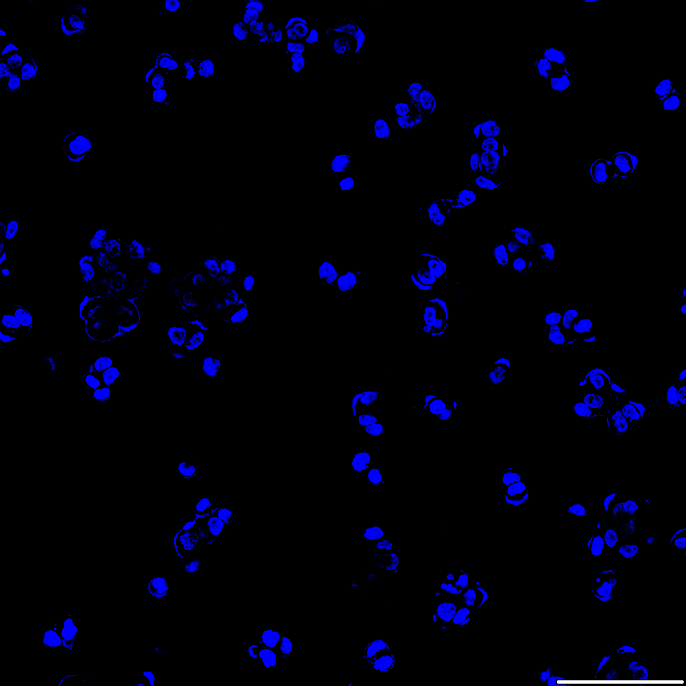

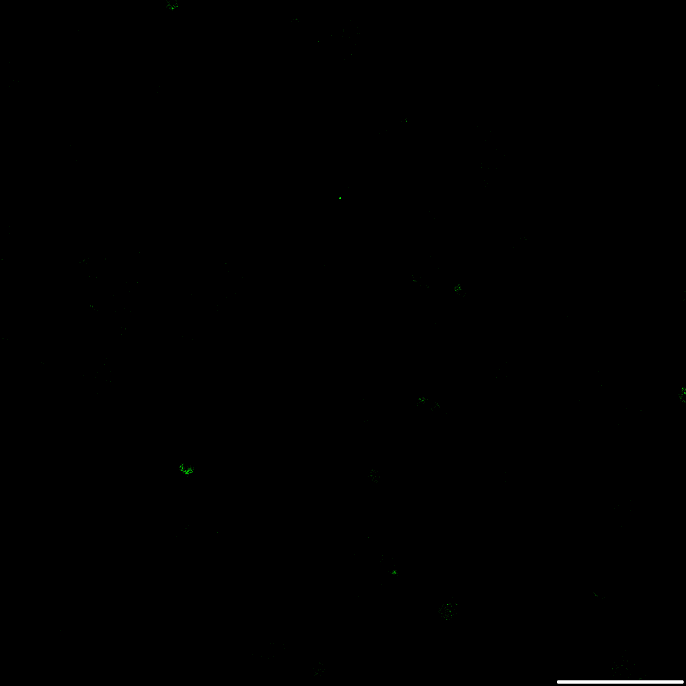

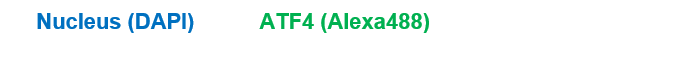

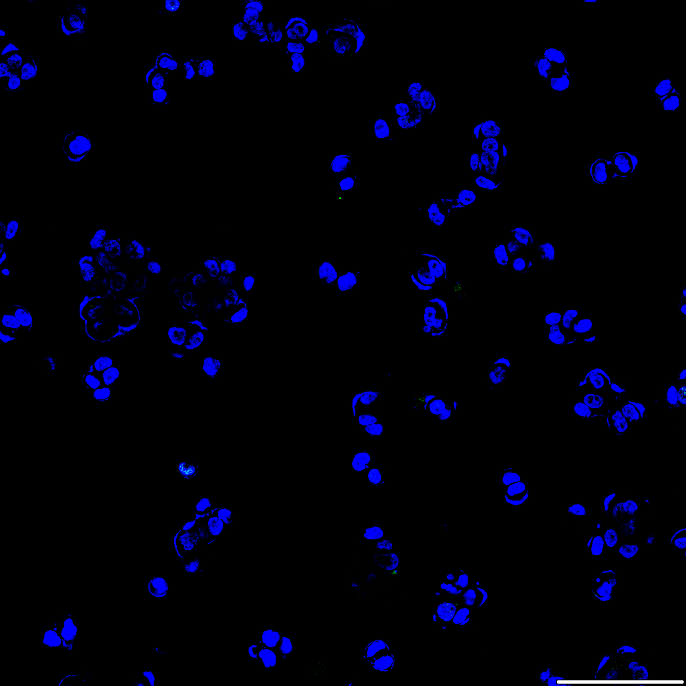

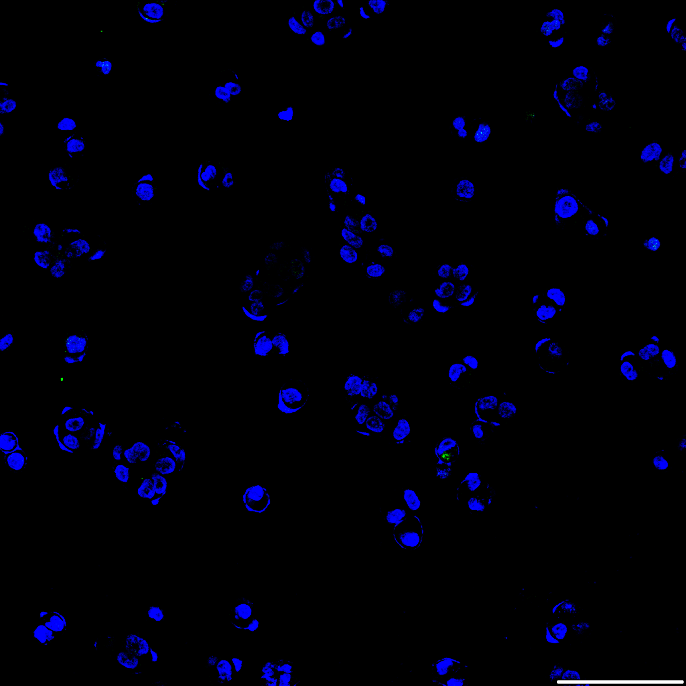

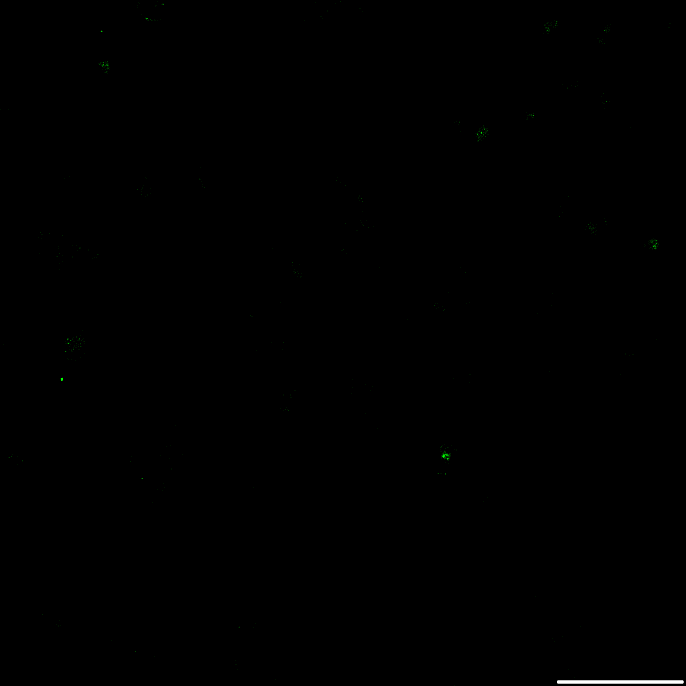

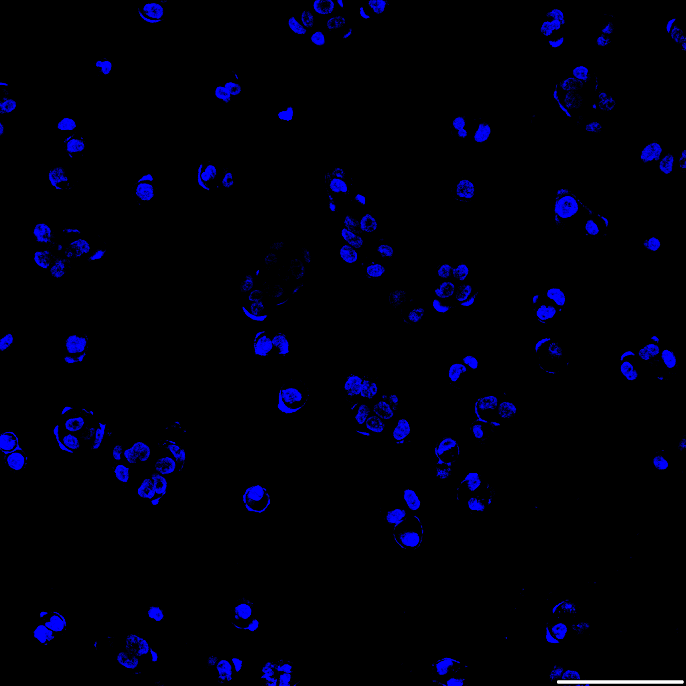

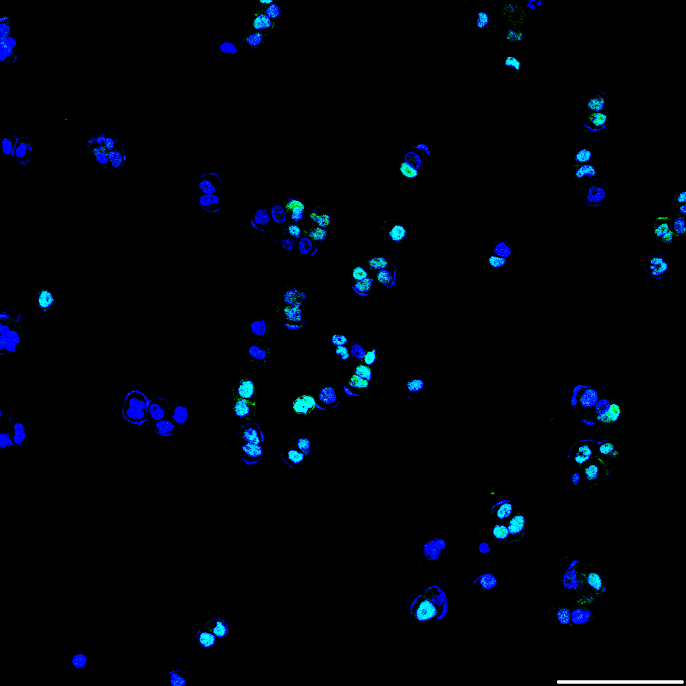

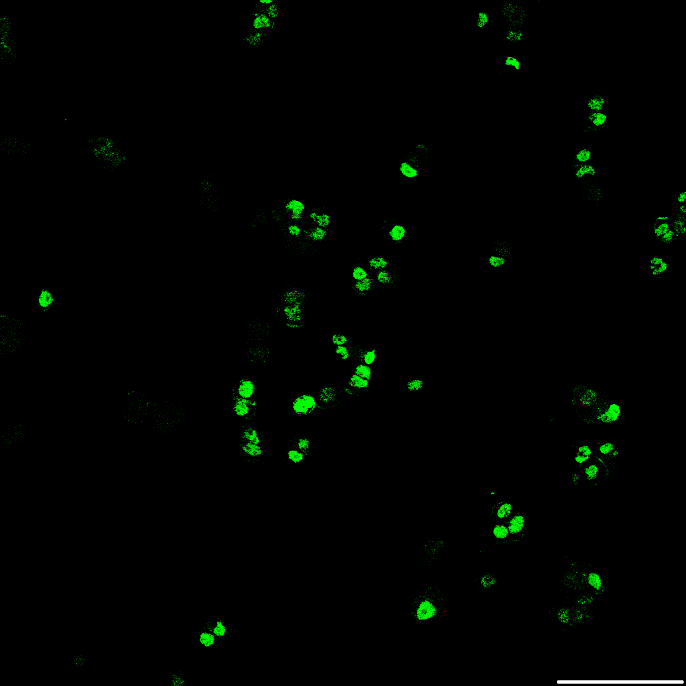

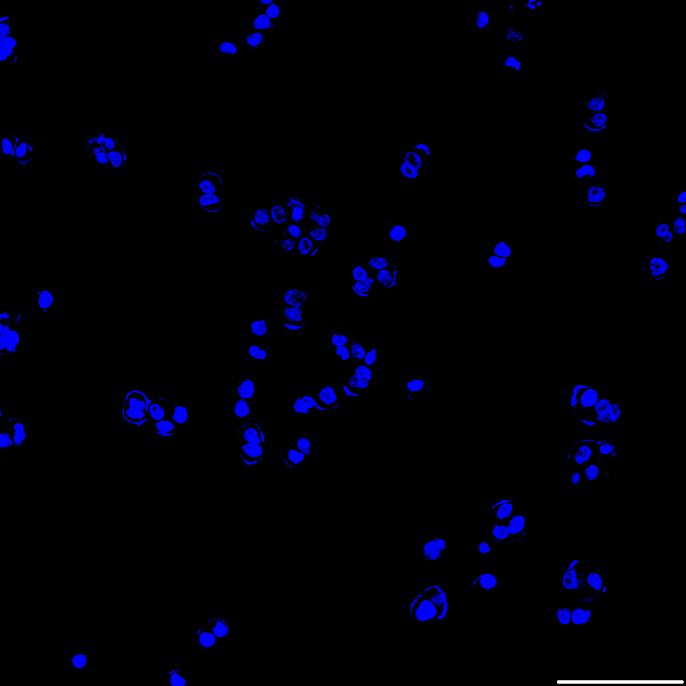

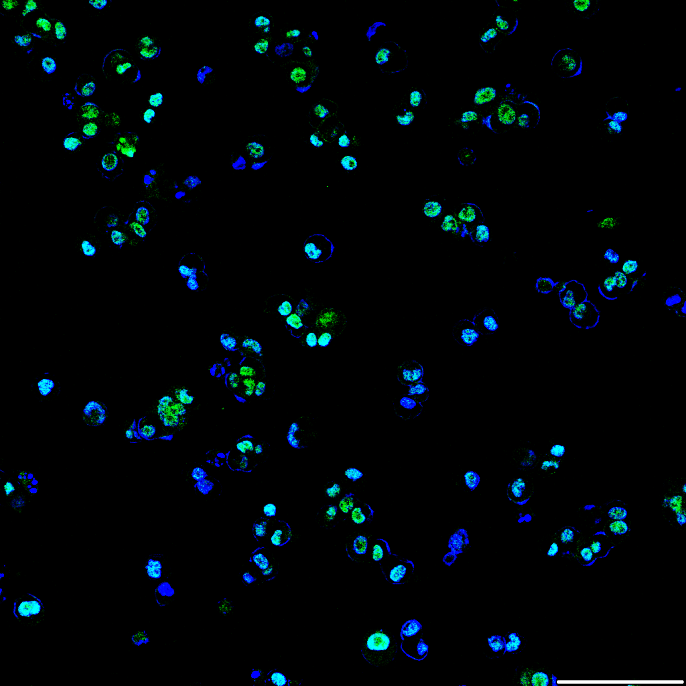

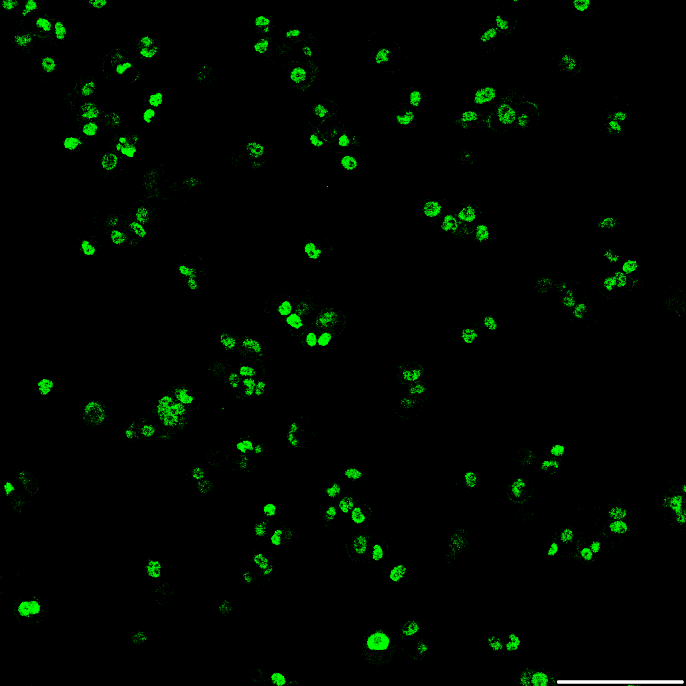

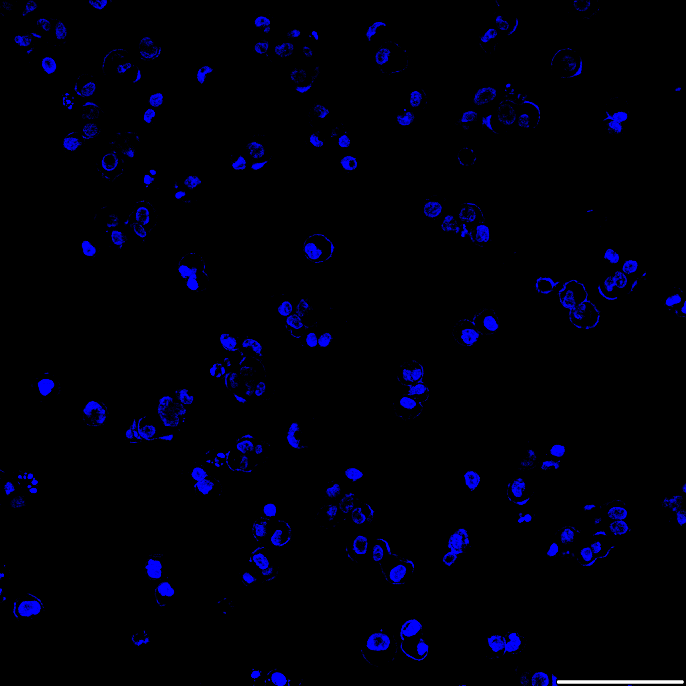

**Supplementary Figure 8. Confocal images showing the upregulation of ATF4 expression by Spiperone**

HepG2.2.15 cells were treated with PBS, TDF or Spiperone (5 μM and 10 μM). Confocal images showing ATF4 (Alexa Fluor 488) in HepG2.2.15 cells. The nuclei were stained with DAPI (blue). The magnification is 20X. Scale bar, 10 μM. TDF, tenofovir disoproxil fumarate.

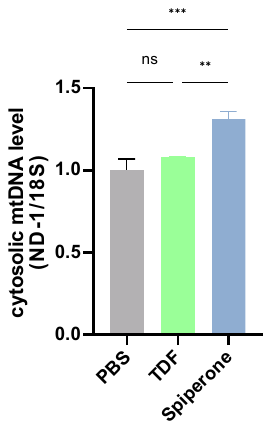

**Supplementary Figure 9. Increase in cytosolic mtDNA (ND-1) release by Spiperone**

HepG2.2.15 cells were treated with PBS or Spiperone for 6h. Cytosolic mtDNA (ND-1) was quantified in the cytosolic fraction, and the 18S rRNA (housekeeping gene) level was measured in the total lysate by qPCR. ND-1 value was normalized to 18S rRNA. Data indicate the mean ± S.D. of three independently performed experiments. * *p* < 0.05, ** *p* < 0.01 and *** *p* < 0.001. qPCR, quantitative polymerase chain reaction.

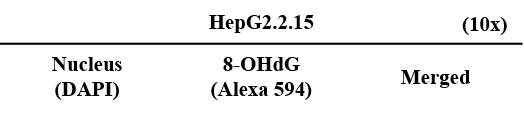

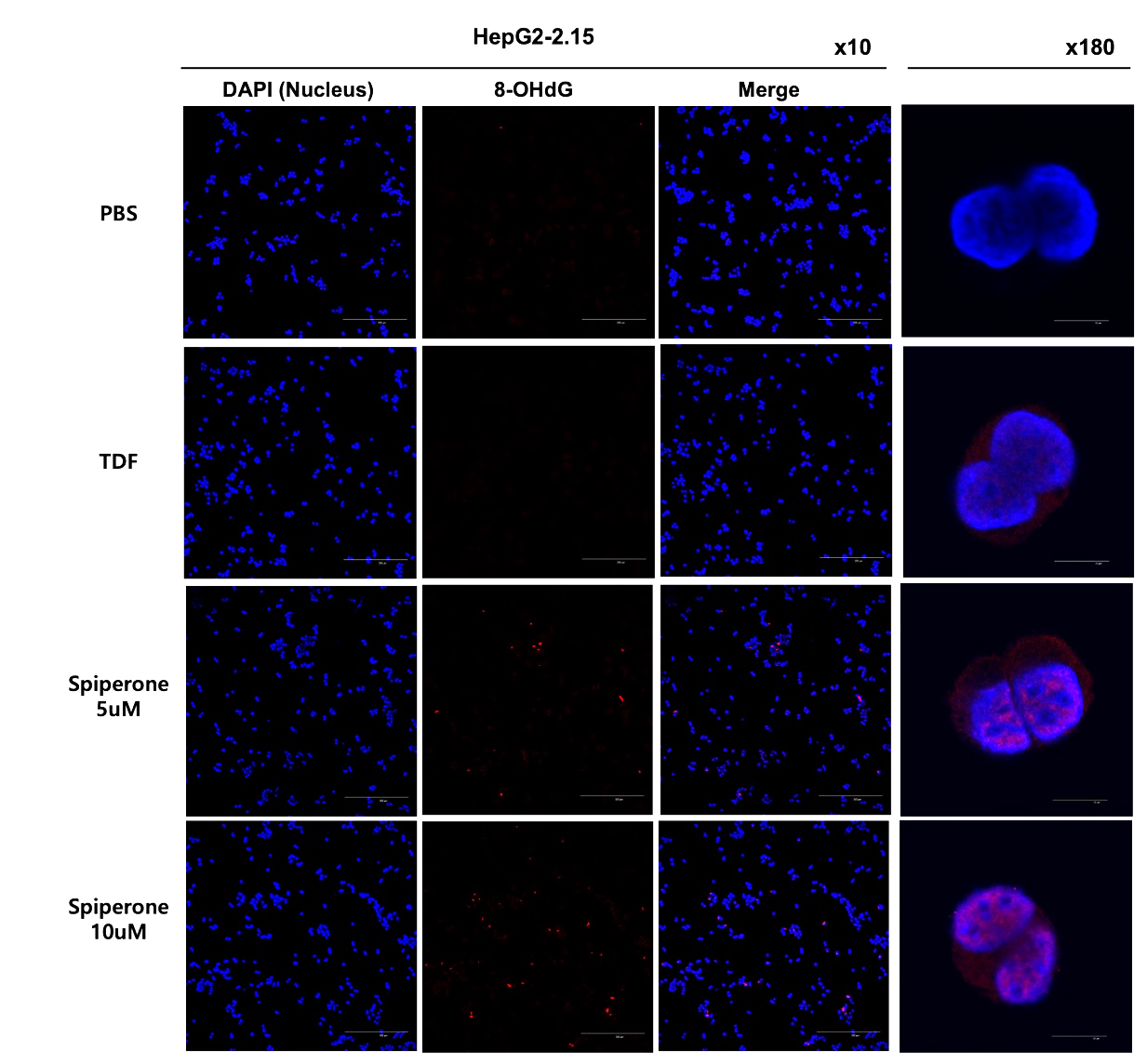

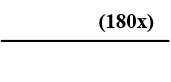

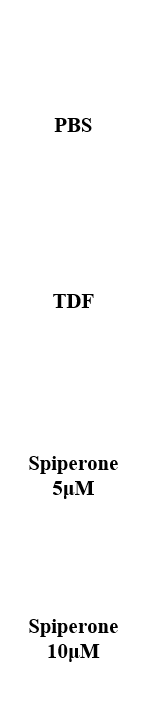

**Supplementary Figure 10. Confocal images showing upregulated 8-OHdG expression by Spiperone**

HepG2.2.15 cells were treated with PBS, TDF or Spiperone (5 μM and 10 μM). Confocal images showing 8-OHdG (Alexa Fluor 594) in HepG2.2.15. The nuclei were stained with DAPI (blue). The magnifications are 10X and 180X. Scale bar, 200 μM. TDF, tenofovir disoproxil fumarate.

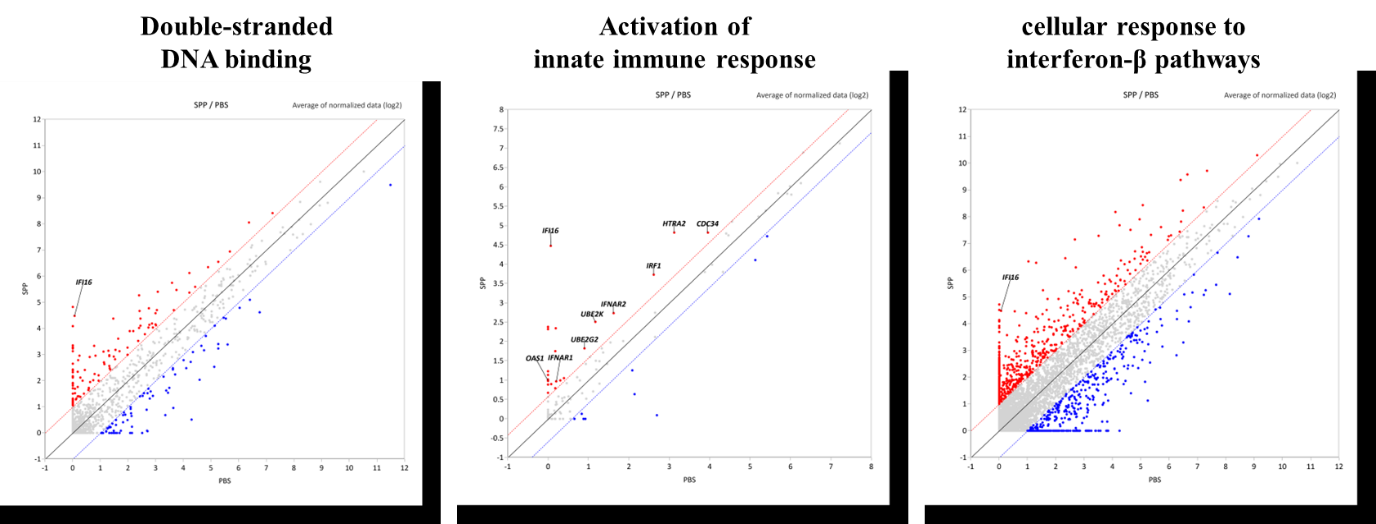

**Supplementary Figure 11. Scatter plots showing gene expression profiles in HepG2.2.15 cells treated with Spiperone compared with PBS control.**

Normalized RNA-seq data (log2 scale) are plotted as average expression values in PBS (x-axis) versus Spiperone (y-axis). Red and blue dots indicate significantly upregulated and downregulated genes, respectively, with gray dots representing genes with nonsignificant changes in expression. Across all three mechanistic pathways analyzed (double-stranded DNA binding, activation of innate immune response, and cellular response to interferon-β), IFI16 was consistently upregulated following spiperone treatment, suggesting a potential central role in spiperone-mediated antiviral responses.

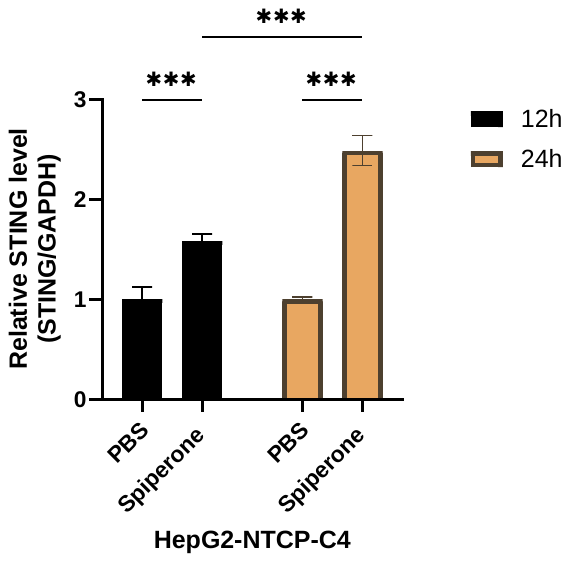

**Supplementary Figure 12. The mRNA level of STING was quantified by qPCR**

HepG2-NTCP-C4 cells were treated with PBS or Spiperone for 12 h and 24 h. The relative mRNA level of STING was measured by qPCR and normalized to GAPDH. Data indicate the mean ± S.D. of three independently performed experiments. * *p* < 0.05, ** *p* < 0.01 and *** *p* < 0.001. qPCR, quantitative polymerase chain reaction.

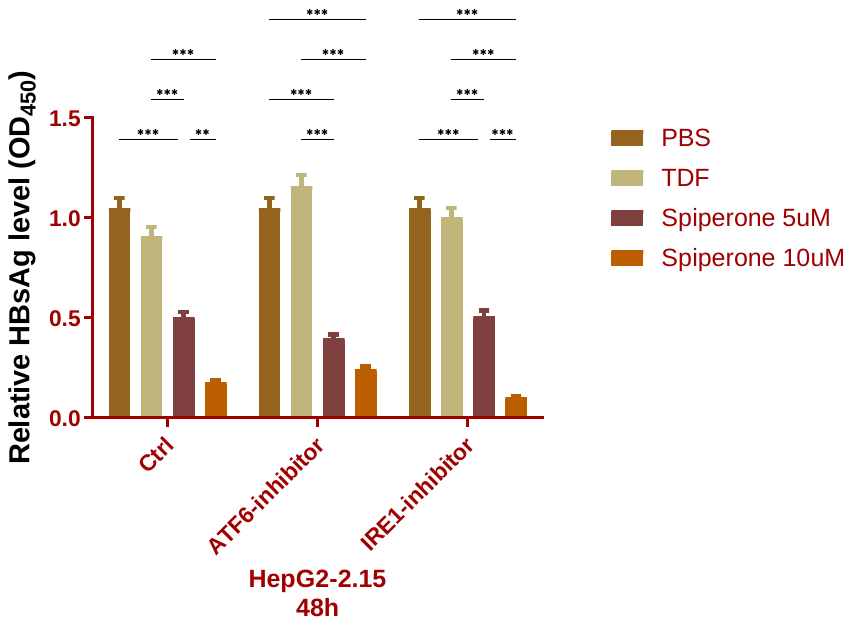

**Supplementary Figure 13. Inhibition of ER stress-induced unfolded protein response mediator**

HepG2.2.15 cells were treated with ATF6-siRNA or IRE1-siRNA. After 48 h of siRNA treatment, the medium was replaced with media containing PBS, TDF or Spiperone. HBsAg in the supernatants were measured by ELISA. Data indicate the mean ± S.D. of three independently performed experiments. * *p* < 0.05, ** *p* < 0.01 and *** *p* < 0.001. HBsAg, HBV surface antigen; TDF, tenofovir disoproxil fumarate.

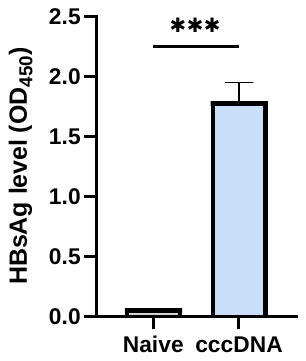

**Supplementary Figure 14. HBsAg levels in a mouse model established by injection of AAV-1.04x virions.**

A model was established in C57BL/6 mice (male) were constructed by injection of AAV-HBV1.04x virions. HBsAg levels in mouse serum samples were measured by ELISA one week after injection to verify the establishment of the cccDNA model. Data indicate the mean ± S.D. of three independently performed experiments. * *p* < 0.05, ** *p* < 0.01 and *** *p* < 0.001. HBV, Hepatitis B virus; HBsAg, HBV surface antigen.

**Supplementary Table S1. List of Primers.**

| **Primer** | **Forward** | **Reverse** |
| --- | --- | --- |
| HBs | 5’-TTG ACA AGA ATC CTC ACA ATA CC-3’ | 5’ -GGA GGT TGG GGA CTG CGA AT- 3’ |
| pgRNA | 5’- GCC TTA GAG TCT CCG GAA CA-3’ | 5’- GAG GGA GTT CTT CTT CTA GG-3’ |
| cccDNA | 5’ - CCG TGT GCA CTT CGC TTC A- 3’ | 5’- GCA CAG CTT GGA GGC TTG A- 3’ |
| cccDNA selective primer | 5’- GGG GCG CAC CTC TCT TTA -3’ | 5’- AGG CAC AGC TTG GAG GC-3’ |
| cccDNA non-selective primer | 5’ -CAC TCT ATG GAA GGC GGG TA -3’ | 5’- TGC TCC AGC TCC TAC CTT GT -3’ |
| Human PERK | 5’- TGG TGG TGC TTC GAG CCA GG -3’ | 5’- AAT GCC TGG GAC GTG GTG GC -3’ |
| Human ATF4 | 5’- GAC CGA AAT GAG CTT CCT GA- 3’ | 5’- ACC CAT GAG GTT TGA AGT GC- 3’ |
| Human ATF6 | 5’- TTC CCT CCT GGT GGA ATT TG -3’ | 5’- AGG CCA CTC TGC TTT CCA AC- 3’ |
| Human XBPs1 | 5’-TGC TGA GTC CGC AGC AGG TG-3’ | 5’-GCT GGC AGG CTC TGG GGA AG-3’ |
| Human STING | 5’- GAG CAG GCC AAA CTC TTC TG-3’ | 5’- TGC CCA CAG TAA CCT CTT CC-3’ |
| Human ND-1 (mitochondrial DNA) | 5′- CGGGCTACTACAACCCTTCG -3’ | 5′- GCGATGGTGAGAGCTAAGGT -3’ |
| Human ND-1 (mitochondrial DNA) | 5’- CTC TTC GTC TGA TCC GTC CT-3’ | 5’- TGA GGT TGA GGT CTG TTA GT-3’ |
| Human ND-2 (mitochondrial DNA) | 5’- GTA GAC AGT CCC ACC CTC AC-3’ | 5’- TTG ATC CCG TTT CGT GCA AG-3’ |
| Human 18S rRNA | 5′- TAGAGGGACAAGTGGCGTTC -3’ | 5′- CGCTGAGCCAGTCAGTGT -3’ |
| Human GAPDH | 5’-CAC ATG GCC TCC AAG GAG TAA -3’ | 5’-GAG GGT CTC TCT CTT CCT CTT GT -3’ |
| P1-P2 primer | 5’-TTT TTC ACC TCT GCC TAA TCA TC-3’ | 5’- AAA AAG TTG CAT GGT GCT GGT G-3’ |

**Supplementary Table S2. List of Antibodies.**

| **Antibody Name** | **Company** |
| --- | --- |
| Anti-HBcAg (WB, IF) | Abcam, ab8637 |
| Anti-HBcAg (IHC) | Millipore Sigma, 216A-15 |
| HBcAg-Alexa488 conjugated | Santa Cruz Biotechnology (Dallas, TX, USA),  sc-23947 AF488 |
| HBsAg (WB, IHC, IF) | Santa Cruz Biotechnology (Dallas, TX, USA), sc-53299 |
| Anti-GAPDH | Invitrogen, MA5-15738-HRP |
| Anti-p-PERK (WB) | Cell Signaling Technology (Danvers, MA, USA), #3179 |
| Anti-p-PERK (IHC) | Santa Cruz Biotechnology (Dallas, TX, USA), sc-32577 |
| Anti-PERK | Cell Signaling Technology (Danvers, MA, USA), #3192 |
| Anti-p-eIF2a | Cell Signaling Technology (Danvers, MA, USA), #3597 |
| Anti-ATF4 (WB) | Cell Signaling Technology (Danvers, MA, USA), #11815 |
| Anti-ATF4 (IF&IHC) | Santa Cruz Biotechnology (Dallas, TX, USA), sc-390063 |
| Anti-8-OHdG | Santa Cruz Biotechnology (Dallas, TX, USA), sc-393870 |
| Anti-IFI16 | Santa Cruz Biotechnology (Dallas, TX, USA), sc-8023 |
| Anti-STING | Cell Signaling Technology (Danvers, MA, USA), #13647 |
| Anti-phospho-TBK1 | Cell Signaling Technology (Danvers, MA, USA), #5483 |
| Anti-TBk1 | Cell Signaling Technology (Danvers, MA, USA), #3504 |
| Anti-phospho-IRF3 | Cell Signaling Technology (Danvers, MA, USA), #37829 |
| Anti-IRF3 | Cell Signaling Technology (Danvers, MA, USA), #4302 |
| Anti-phospho-NF-κB p65 | Cell Signaling Technology (Danvers, MA, USA), #3033 |
| Anti- NF-κB p65 | Cell Signaling Technology (Danvers, MA, USA), #6956 |
| Anti-IRF9 | Cell Signaling Technology (Danvers, MA, USA), #76684 |
| Anti-phospho-STAT1 | Cell Signaling Technology (Danvers, MA, USA), #9167 |
| Anti-STAT1 | Cell Signaling Technology (Danvers, MA, USA), #9172 |
| Acetyl-Histone H4 (Lys5) antibody | Cell Signaling Technology (Danvers, MA, USA), #8647 |
| Histone H3 (acetyl K27) antibody | Abcam, ab4729 |
| Acetyl-Histone H3 Antibody | Merck Millipore, #06-599 |
| Acetyl-Histone H4 Antibody | Merck Millipore, #06-598 |
| Anti-mouse-Alexa Fluor™ 488 | Invitrogen, #A-11001 |
| Anti-mouse-Alexa Fluor™ 594 | Invitrogen, #A-11005 |
